# HIV-2 Immature Particle Morphology Provides Insights into Gag Lattice Stability and Virus Maturation

**DOI:** 10.1101/2022.02.01.478508

**Authors:** Nathaniel Talledge, Huixin Yang, Ke Shi, Raffaele Coray, Guichuan Yu, William G. Arndt, Shuyu Meng, Gloria C. Baxter, Luiza M. Mendonça, Daniel Castaño-Díez, Hideki Aihara, Louis M. Mansky, Wei Zhang

## Abstract

Retrovirus immature particle morphology consists of a membrane enclosed, pleomorphic, spherical and incomplete lattice of Gag hexamers. Previously, we demonstrated that human immunodeficiency virus type 2 (HIV-2) immature particles possess a distinct and extensive Gag lattice morphology. To better understand the nature of the continuously curved hexagonal Gag lattice, we have used single particle cryo-electron microscopy with a retrovirus to determine the HIV-2 Gag lattice structure for immature virions. The reconstruction map at 5.5 Å resolution revealed a stable, wineglass-shaped Gag hexamer structure with structural features consistent with other lentiviral immature Gag structures. Cryo-electron tomography provided evidence for nearly complete ordered Gag lattice structures in HIV-2 immature particles. We also solved a 1.98 Å resolution crystal structure of the carboxyl-terminal domain (CTD) of the HIV-2 capsid (CA) protein that identified a structured helix 12 supported via an interaction of helix 10 in the absence of the SP1 region of Gag. Residues at the helix 10-12 interface proved critical in maintaining HIV-2 particle release and infectivity. Taken together, our findings provide the first 3D organization of HIV-2 immature Gag lattice and important insights into both HIV Gag lattice stabilization and virus maturation.

## Introduction

Human immunodeficiency virus type 1 (HIV-1) and HIV type 2 (HIV-2) are both etiological agents of acquired immunodeficiency syndrome (AIDS). The HIVs are members of the lentivirus genus of *Retroviridae* [1]. Relative to that of HIV-1, HIV-2 has an attenuated disease phenotype, with lower viral loads in infected individuals, as well as lower rates of both vertical and sexual transmissibility, and a slower progression to AIDS [2–7]. The HIV-2 genome structure is like that of the HIV-1 genome; however, HIV-2 HIV-2 encodes for *vpx* instead of *vpu* [8–10]. HIV-1 emerged in the human population via spillover of simian immunodeficiency virus from chimpanzees or gorillas (SIVcpz, SIVgor), while HIV-2 emerged in humans following spillover of SIV from sooty mangabeys (SIVsm) [11]. HIV-1 and HIV-2 share approximately 60% amino acid identity between Gag polyproteins and just 48% identity at the viral genome level [12]. As lentiviruses, HIV-1 and HIV-2 mature particles possess similar morphologies that include a conically shaped core structure [13, 14]. Immature HIV-1 and HIV-2 particles have been noted to have distinct morphological features [15].

HIV particle assembly is known to occur at the plasma membrane of infected host cells, where multimerization of the Gag structural polyprotein drives the assembly and release of virus particles (for a review, see [16]). The Gag polyprotein consists of three major protein domains: the matrix (MA) domain (which binds membrane), the capsid (CA) domain (which encodes key residues involved in Gag-Gag and CA-CA interactions), and the nucleocapsid (NC) domain (which encodes the key determinants for genomic RNA packaging) [16, 17]. The HIV-1 and HIV-2 Gag polyproteins both encode for two spacer peptides (*i.e.,* SP1 and SP2) and the p6 domain. During virus maturation, the Gag polyprotein is cleaved by the virally encoded protease to produce the mature viral structural proteins. The emergence of these structural proteins leads to formation of a conically shaped CA core.

HIV-1 CA is composed of two α-helical enriched domains connected by a flexible linker [18–20]. The amino-terminal domain (*i.e.,* CA_NTD_) is composed of 7 α-helices (H1 to H7) and includes functional domains that bind to cellular factors such as cyclophilin A (CypA) [18] and TRIM5α [21, 22]. The carboxy-terminal domain (*i.e.,* CA_CTD_) contains 4 α-helices (H8 to H11). The function of the CA_CTD_ is primarily to form inter-molecular contacts that are important to the stability of immature Gag lattice and for the formation of a mature CA lattice that is necessary for particle infectivity.

Multimerization of Gag in immature HIV-1 particles results in hexagonal lattice structure due to extensive interactions among the CA proteins [23–29]. The CA hexamers adopt a wineglass-like structure to accommodate the spherical shape of the viral membrane. The “cup” of the wineglass is composed of six copies of CA_NTD_ on the rim and six copies of CA_CTD_ at the base of the cup. The “stem” of the wineglass is a six-helix bundle (6HB) structure that consists of the C-terminal tail of CA and the SP1 segment (CA-SP1) [30]. The CA hexamer in HIV-1 is further stabilized by the cellular metabolite inositol hexakisphosphate (IP6) that is closely positioned within the positively charged rings of lysine residues in the C-terminal end of the CA_CTD_ (*i.e.,* Gag residues K290 and K359, or CA residues K158 and K227) [30, 31]. The Gag lattice is stabilized by the molecular interactions of helix 1 and 2 (*i.e.,* H1 and H2) in the CA_NTD_ at local two-fold and three-fold axes, respectively, and the CA_CTD_ interactions (H9) at the local two-fold axes [30].

The HIV-1 capsid core has a fullerene structure composed of approximately 250 CA hexamers and 12 pentamers [32–37]. In contrast to the CA hexamers in the immature lattice, each capsomer in the mature CA lattice adopts a bell-shaped structure. The CA_NTD_ forms the inner ring and the closed end of the bell. The CA_CTD_ forms the outer ring connecting the neighboring capsomers. The HIV-1 CA_CTD_ H10 forms the trimer interface and H9 forms the dimer interface between the adjacent CA molecules.

Limited structural details are available for the HIV-2 Gag protein, with the crystal structure of the HIV-2 CA_NTD_ being the only high-resolution protein structure of HIV-2 CA solved to date (PDB ID: 2WLV) [38]. A previous comparative analysis of immature retrovirus particle morphologies by cryo-electron microscopy (cryo-EM) revealed that HIV-2 immature particles possessed distinct differences from that of HIV-1 or other immature retrovirus particles analyzed [15]. In contrast to what was observed for HIV-1, immature HIV-2 particles had a relatively narrow range of particle diameters and possessed nearly complete Gag lattice structures as assessed by qualitative measurements of the particle morphology. However, to date, no high-resolution structure of immature particles has been analyzed for HIV-2.

To help address this knowledge gap, and to help define the basis for the morphological differences between HIV-1 and HIV-2, we conducted cryo-EM and cryo-electron tomography (cryo-ET) of HIV-2 particles and confirmed that HIV-2 immature particles have a high occupancy of immature Gag lattice underneath the viral membrane. We determined the HIV-2 Gag hexamer structure at 5.5 Å resolution by using a cryo-EM single particle reconstruction (SPR) method. Furthermore, we also solved a 1.98 Å crystal structure of the HIV-2 CA_CTD_, which revealed a novel, extended 3_10_ helix at the CTD of CA (H12). Residues that are involved in H10 and H12 interactions in the crystal dimer were found to be critical for maintaining HIV-2 particle infectivity. Taken together, our observations demonstrate – for the first time – that the HIV-2 Gag lattice possesses few interspersed gaps relative to that of HIV-1 [24], with novel findings of critical amino acid residues within the HIV-2 CA_CTD_ that are required for infectious particle production. These findings provide insights into the morphological differences observed between HIV-1 and HIV-2 immature particles, as well as general implications for HIV Gag lattice stabilization and virus maturation.

## Results

### Cryo-EM reconstruction of the immature HIV-2 Gag lattice

We computed the reconstruction map of HIV-2 Gag lattice by using a cryo-EM SPR method **(Figure 1A, S1)**. The boxed particle centers locate at the center of the lattice region of the imaged immature particles. This region represents the side views of the segmented Gag lattice, allowing for rapid analysis of heterogeneous sized viral particles. Using the six-fold rotational symmetry from the Gag hexamer we computed a reconstruction of the underlying capsid lattice (**Figure 1B-D**). The final reconstruction of the HIV-2 CA lattice in immature particles was resolved to 5.5 Å resolution that contains 19 hexamer units of the HIV-2 Gag lattice **(Figure 1C-D, S2)**.

**Figure 1.**
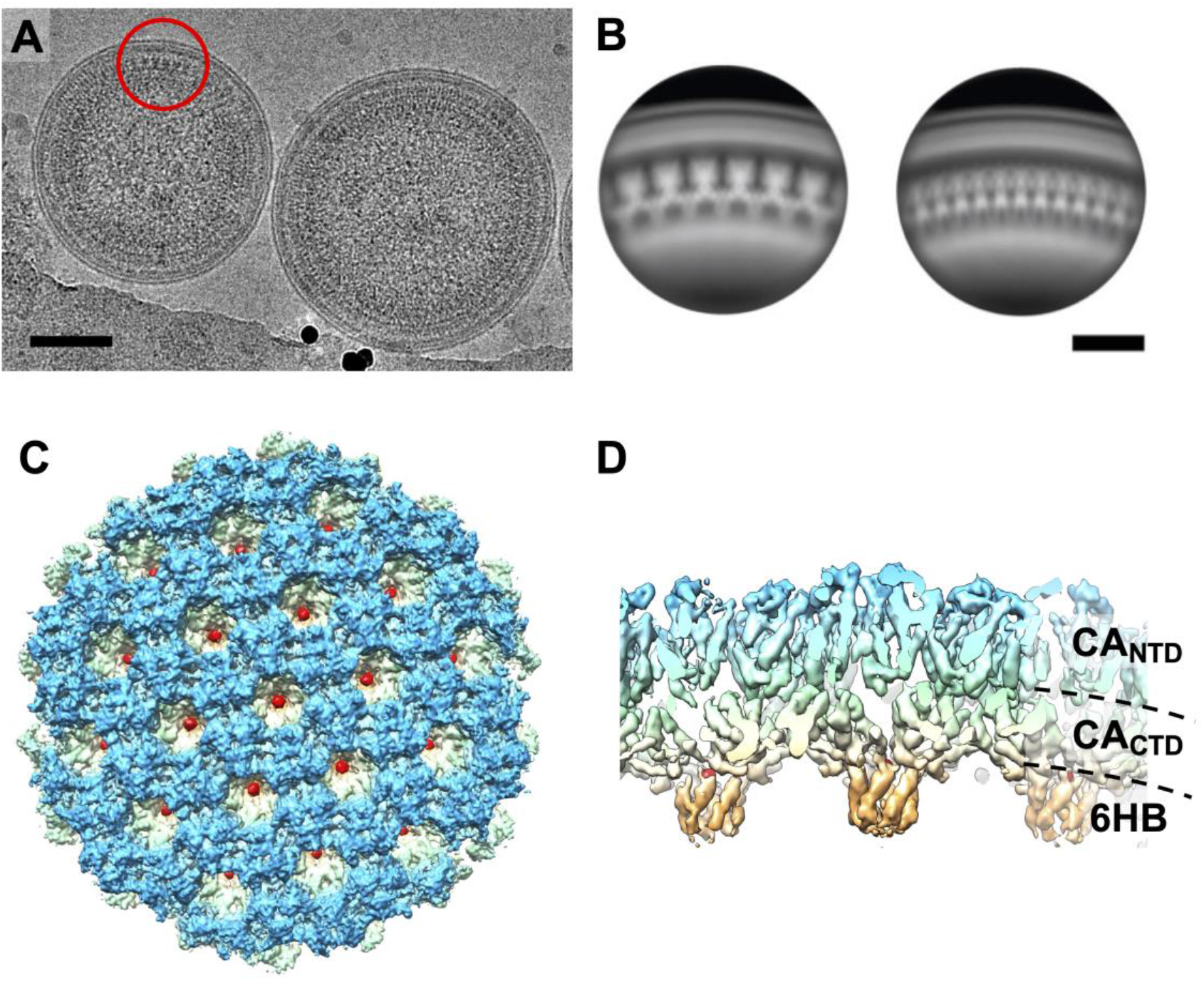
HIV-2 Gag lattice structure. (A) Cryo-EM micrograph of immature HIV-2 particles. The red circle indicates an example area of extracted particles for the single particle reconstruction procedure. (B) Representative 2D class averages of the resulting particle extraction at the tangent of the immature HIV-2 particles. The scale bars represent 50 nm. (C)Top and (D) side views of the 5.5 Å reconstruction map showing densities of CA_NTD_ (blue), CA_CTD_ (light green and yellow), CA-SP1 six helix bundle (6HB, orange), and IP6 (red).

### Crystal structure of HIV-2 CA_CTD_ has a distinct C-terminal 3_10_ helix

To help elucidate the molecular basis of the HIV-2 Gag lattice interactions, we expressed and purified the CA CTD residues 145-230 (HIV-2 CA_CTD_, Gag residues T280 to M365) to homogeneity for structure determination. HIV-2 ROD CA_CTD_ shares 69% sequence identity with that of HIV-1 NL4-3. This subdomain of HIV-2 CA structure has not been previously reported. The purified protein was crystallized, and the structure determined to 1.98 Å resolution (**Figure 2, Table S1**). The crystal belonged to the space group C2, with a monomer of HIV-2 CA_CTD_ in the asymmetric unit. Two HIV-2 CA_CTD_ molecules form a symmetric dimer with the 2-fold axis coinciding with the crystallographic dyad. We were able to model CA residues T147 to M230 (Gag residues T282 to M365).

**Figure 2.**
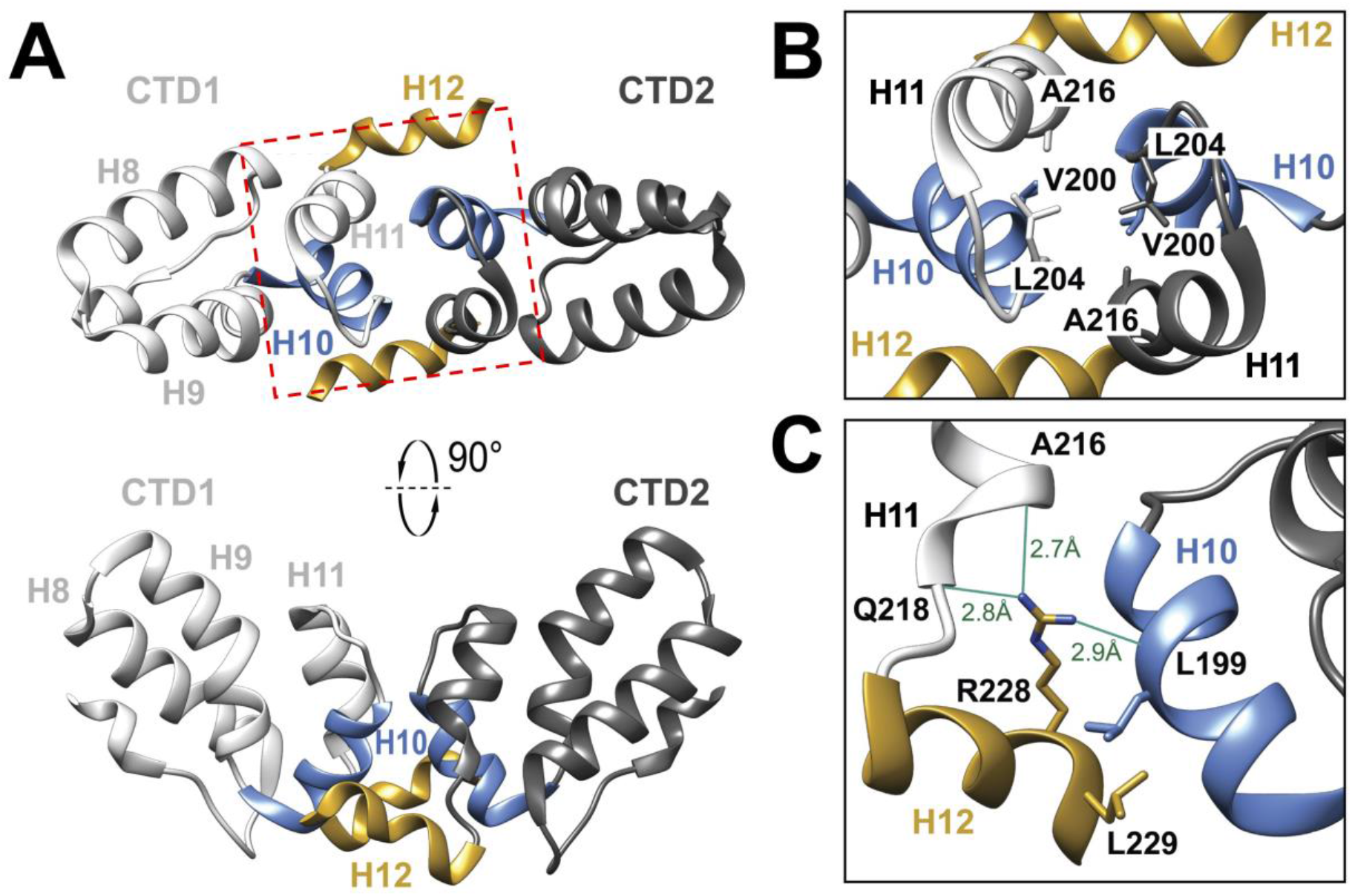
HIV-2 CA_CTD_ crystal Structure. (A) The structure of HIV-2 CA_CTD_ dimer from CA residues T147-M230. H8, H9, H10, and H11 represent the four α-helices in HIV-2 CA_CTD_. H12 is a 3_10_ helix at the C-terminus of the molecule. CTD1 and CTD2 represent two molecules in the dimer. The red box marks the H10-H11 dimer interface which is delineated in B. (B) Magnified view of the H10-H11 crystallographic dimer interface highlighting the residues V200, L204, and A216. (C) Residues L199, R228, and L229 are located at the HIV-2 CA_CTD_ H10-H12 crystallographic dimer interface. The guanidinium group of CTD2-R228 forms hydrogen bonds with the backbone oxygen atoms of CTD1-L199, CTD2-A216, and CTD2-Q218.

Like that of other lentiviral CA_CTD_ domains [38–41], the HIV-2 CA_CTD_ is primarily α-helical. Furthermore, its tertiary structures (the helix 11, H11, and its upstream domain) superimpose with that HIV-1 CA_CTD_ (PDB ID: 1A43) [41] with backbone RMSD of 0.74 Å. The residues downstream of H11 form an extended C-terminal 3_10_ helix (referred to as helix 12, or H12).

The HIV-2 CA_CTD_ dimer interface involves H10 in one monomer contacting with H10, H11 and H12 in another monomer (**Figure 2B-C**). In contrast, the crystallographic CTD dimer interface in HIV-1 CA_CTD_ is composed of the intermolecular interactions between H9 of each monomer [41]. Specifically, the unique HIV-2 CA_CTD_ dimeric interface is supported by hydrogen bonds with the backbone oxygen atoms of CTD1-L199 (H10), CTD2-A216 (H11), CTD2-Q218 (H11) and the guanidinium group of CTD2-R228 (H12) **(Figure 2C)**. The interface mediated by intermolecular contacts between H10 and H12 contrasts with other retroviruses CA_CTD_ as this portion of the CTD is disordered in the absence of the SP1 peptide in other retroviral CA_CTD_ structures [38, 41].

### The HIV-2 hexagonal Gag lattice resembles that observed in HIV-1

The 5.5 Å reconstruction map of the immature HIV-2 Gag lattice allowed for modeling of the Gag residues S150 to I373 (CA residues S15 to M230 and SP1 residues A1 to I8) into the electron density (**Figure 3**). The overall molecular fitting of our structure indicates a similar structural conformation to that of HIV-1. For example, six copies of the CA and SP1 domains are organized into a wineglass-shaped structure, with CA proteins representing the cup and the CA-SP1 6HB representing the stem **(Figure 3 A-B)**. The NTD and CTD domains of the HIV-2 CA twist in a left-hand manner, while the 6HB twists in a right-hand manner **(Figure 3C)**. The length of the 6HB density layers is ∼25 Å, which can accommodate the predicted 16 amino acid residues of the CA-SP1 region (*i.e.,* Gag residues P358 to I373). The extra globular-shaped densities observed at the top of the 6HB are attributable to IP6 **(Figure 3A-B)**. This observation is consistent with what has been previously reported for the HIV-1 immature Gag lattice structure [30, 42], and is consistent with the finding that IP6 plays a role in HIV-2 immature particle assembly [31]. Like HIV-1, the HIV-2 Gag inter-hexamer interface is also maintained by the molecular contact regions at both the two-fold and three-fold symmetric axes (**Figure 3D-F**). A number of amino acid residues at the interaction interface are conserved between HIV-1 and HIV-2. For example, the HIV-2 CA H9 residues CA W183 and M184 (Gag residues W318 and M319) are located at the two-fold interface of the Gag lattice. Such structural similarity among lentiviruses indicates the conserved stability of the Gag hexamer as a structural building block of the immature Gag lattice.

**Figure 3.**
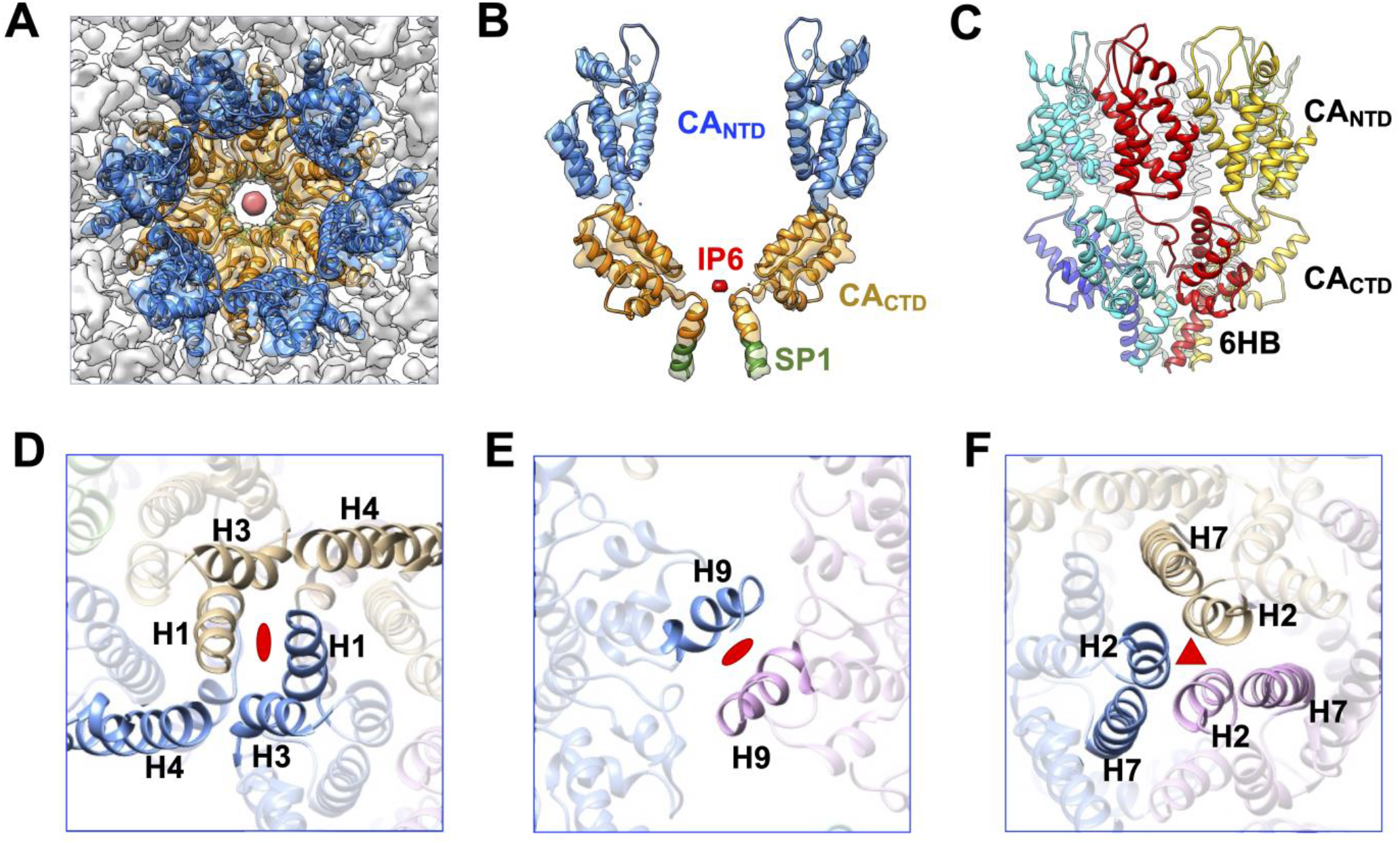
Model of HIV-2 CA hexamer and lattice interfaces. (A) Top view of the central hexamer reconstruction density fitted with the HIV-2 CA model (PDB ID: 8FZC). Blue: CA_NTD_; Orange: CA_CTD_; Red: IP6. (B) Side view of two capsid monomers zoned out of the central hexamer. Green: SP1. (C) Side view of HIV-2 hexamer model color coded by the CA proteins. The depiction shows the left-hand twist of the CA proteins and the right-hand twist of the six-helix bundle (6HB) formed by CA-SP1. (D) View of the NTD two-fold interface in the Gag lattice involving H1. (E) View of the CTD two-fold interface formed by H9. (F) View of the NTD three-fold interface formed by H2.

### Mutations of H10 and H12 residues decrease HIV-2 particle infectivity

To determine if the extended HIV-2 H12 revealed in the crystal structure has impact on aspects of the retroviral life cycle we made disruptive mutations at critical residues at the HIV-2 CA_CTD_ H10 and H12 crystal interface **(Figure 4, Figure S7)**. Specifically, residues L199, R228, and L229 observed at the CA_CTD_ crystal structure interface (**Figure 2C**) were mutated to introduce either modest disruptions (L199G, R228A, and L229G) or strong disruptions (L199K, R228E, and L229K). To assess particle assembly defects, we analyzed particle production from expressing a codon-optimized HIV-2 Gag construct harboring the individual mutations in HEK293T/17 cells. Viral supernatants were assessed via immunoblot for the relative amount of Gag present to WT HIV-2 Gag (**Figure 4A; Figure S9**). Two mutation sites in H12 R228 and L229 showed a significant decrease in produced immature particles where the H10 L199G and L199K point mutations showed no significance difference to that of WT Gag levels.

**Figure 4.**
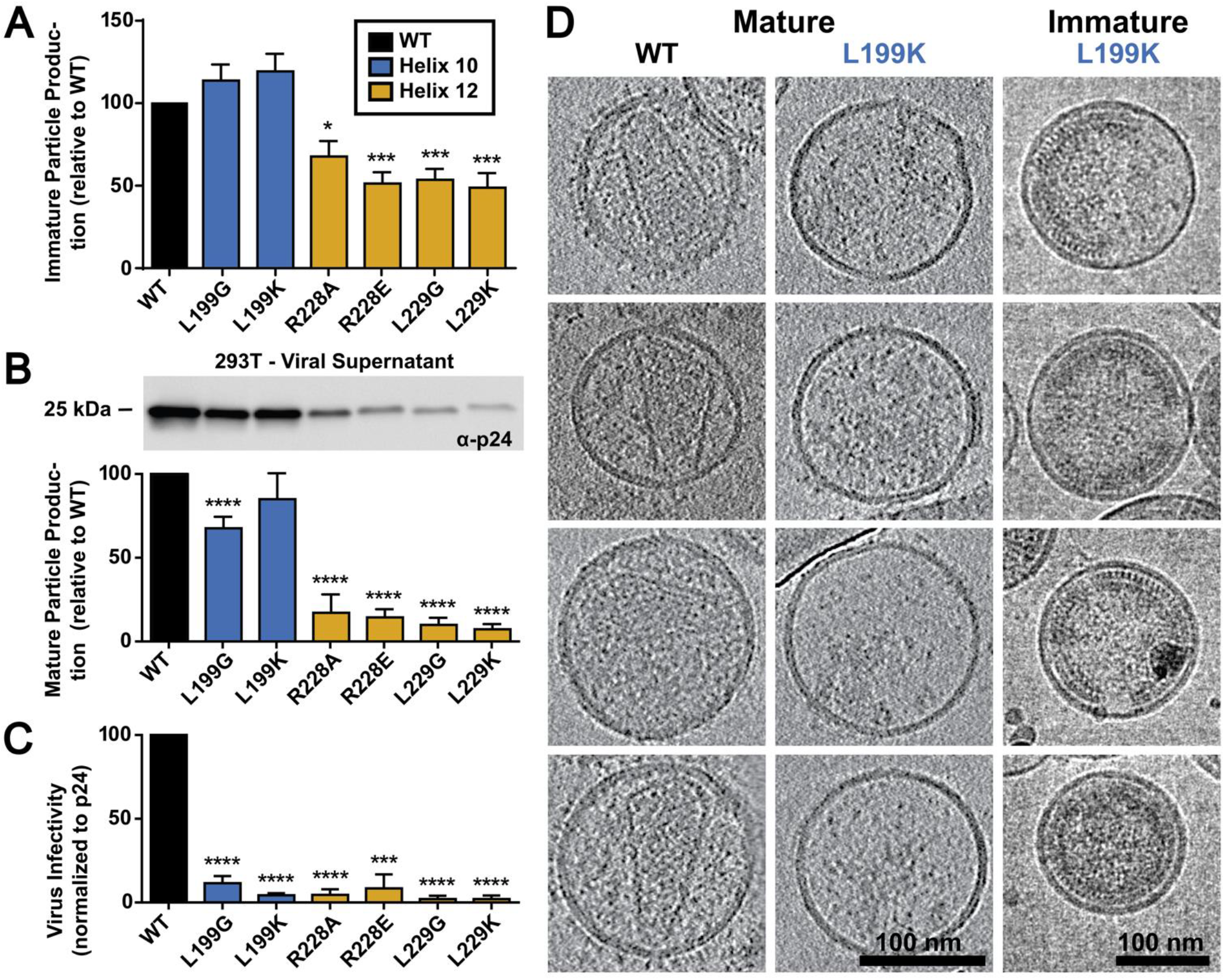
Analysis of mutations in HIV-2 CA_CTD_ H10 and H12 on particle production and infectivity. (A) The effect of H10 and H12 mutations on immature particle production, (B) mature particle production including an immunoblot of the relative levels of CA (p24) present in the viral supernatant, and (C) on particle infectivity is shown. Results are from three independent experiments: average value ± standard error of the mean shown. Significance relative to WT as assessed by using an unpaired t-test. ****, P < 0.0001; ***, P < 0.001; **, P < 0.01; *, P < 0.05. (D) Morphology of mature and immature HIV-2 WT and L199K mutant particles. The left and the middle columns are representative slices of cryo-ET reconstructions of the HIV-2 MIG vector, and its mutant (L199K). The right column shows cryo-EM images of the immature particles for the L199K mutant.

To determine if these mutations influenced mature particle morphology, an HIV-2 ROD10-based vector (*i.e.,* an *env*-minus HIV-2 vector containing the mCherry-IRES-GFP reporter gene expression cassette) was co-expressed with VSV-G to produce single-cycle mature infectious particles in HEK293T/17 cells [39]. These particles were used to assess mature particle production and morphology (**Figure 4B; Figure S10**). All of the H12 mutations significantly decreased (i.e., ∼10 fold) particle production compared to that of WT. The H10 mutation led to a significant decrease for the L199G (i.e., ∼60% that of WT), but the L199K mutation showed no significant difference (**Figure 4B; Figure S10**).

The HIV-2 vector virus particles that harbor dual fluorescent reporters were then used to assess particle infectivity by infecting MAGI cells with equivalent amounts of virus as assessed by CA levels in the produced viral supernatant (**Figure 4B; Figure S10**). Specifically, the H10 or H12 mutants harboring particles showed significant defects in infectivity as compared to a WT infection. Particle production was ∼10 to 20-fold lower to that of WT HIV-2 particles (**Figure 4C**).

Two patterns emerged when assessing mature particle production (**Figure 4B; Figure S10**) and particle infectivity (**Figure 4C**) for the HIV-2 H10/H12 mutants. The H12 residues all severe defects in particle production and infectivity. Intriguingly, mutation of the L199 position in H10 retained particle efficient particle production, but significantly reduced particle infectivity. Both immature and mature HIV-2 CA L199K particles were visualized by cryo-ET to assess particle morphology (**Figure 4D**). HIV-2 WT particles had a canonical mature core morphology with the presence of a fullerene cone in a subset of the produced particles. The HIV-2 L199K immature particles had intact and extensive Gag lattice below the viral membrane, but the L199K MIG mutant had no particles with intact capsid cores, indicating a defect in mature particle formation. To rule out the possibility that mutations in CA impacted the recognition by anti-CA antibodies, a control immunoblot was done in which anti-MA antibodies were used for Gag detection. Immature particle production (**Figure S11**) was comparable to that observed with the anti-CA antibodies (**Figure S9**).

### Cryo-ET reveals high Gag occupancy and minimal gaps in the HIV-2 Gag lattice

The SPR map of immature HIV-2 particles revealed an ordered Gag lattice morphology that occupies an extensively large region underneath the viral membrane, consistent with our group’s previous reports when analyzing the cryo-EM images [15]. To fully characterize this observation, cryo-ET was conducted on immature HIV-2 particles purified from HEK 293T/17 cells. Five cryo-ET tilt series of HIV-2 immature particles were reconstructed in IMOD. Then subtomogram averaging (StA) workflows were implemented in Dynamo followed by RELION-4.0 (**Figure 5, S3**) [43].

**Figure 5.**
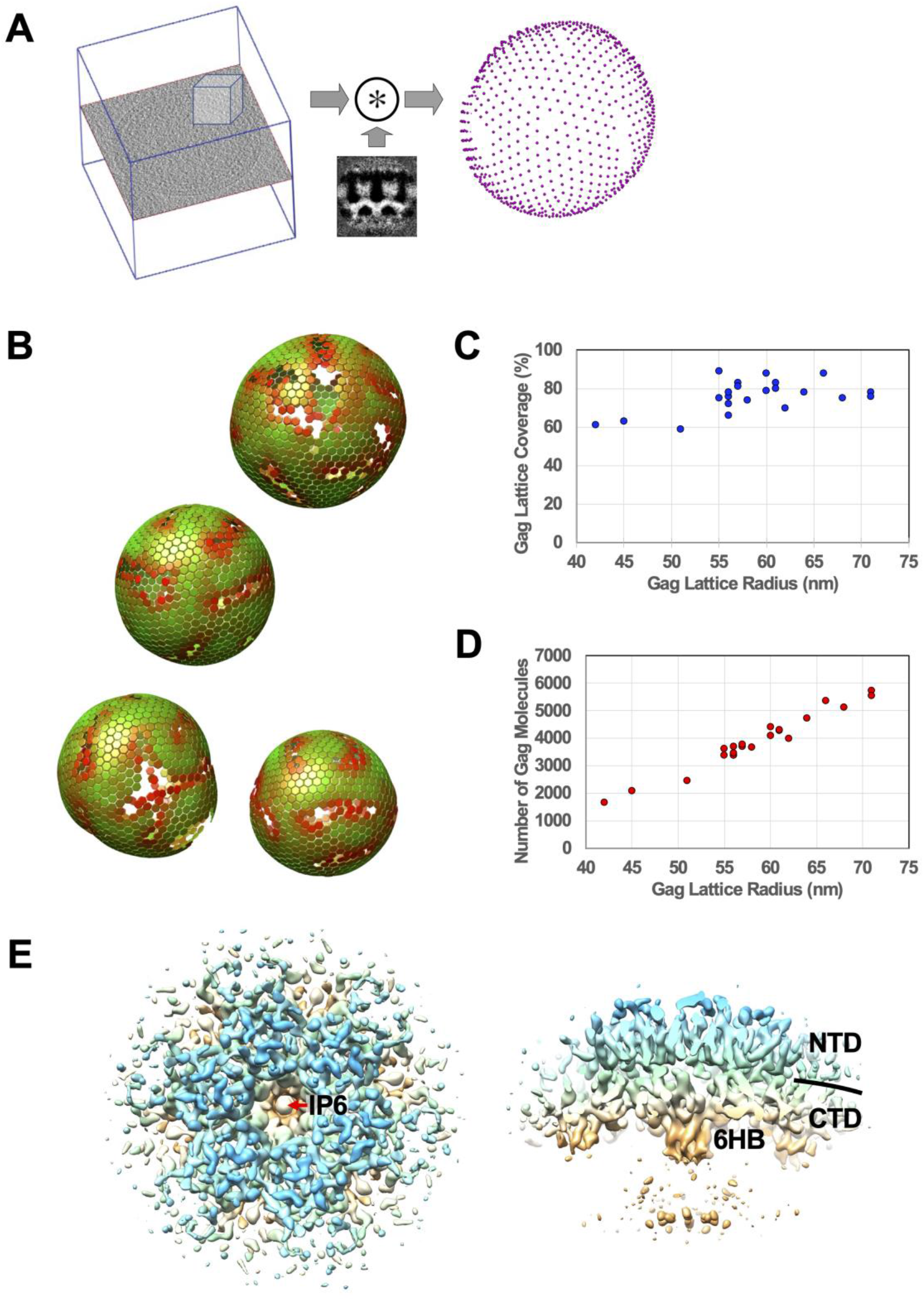
Cryo-ET and subtomogram averaging (StA) analysis of HIV-2 immature particles. (A) Illustration of convolution calculation between a region cropped from a particle (left) with the StA average of the Gag hexamer (middle). The calculation produces the Gag hexamer localizations (right). (B) A graphical depiction of the hexagonal Gag lattices, determined by Dynamo, for four HIV-2 immature particles within the same tomogram. The color denotes how well that hexamer matches with the StA model, with green having a higher correlation coefficient score than red. (C) Gag lattice coverage sorted by the estimated Gag lattice radius per particle. The radius of the Gag lattice is defined as the radius of the linker region (between CA_NTD_ and CA_CTD_) of the CA density layer. (D) Estimate of the total number of Gag molecules underneath the viral membrane sorted by the estimated Gag lattice radius per particle. (E) The top and side 3D rendering views of the StA average of a Gag hexamer at 9.1 Å resolution. IP6: inositol hexakisphosphate; 6HB: six helix bundle; NTD: CA_NTD_; CTD: CA_CTD_.

Gag hexamer lattice positioning was determined in Dynamo by an iterative convolution procedure that correlates the sub-volumes cropped from the IMOD reconstruction maps at regions corresponding to the Gag lattice with the StA averaged map centered at the Gag hexamer model **(Figure 5A)**. The calculated HIV-2 Gag hexamer positions within the viral particles were found to have an almost complete hexagonal lattice pattern with small, interspersed gaps or crevice regions indicated by low positional cross-correlations from our analysis (**Figure 5B-C**, **Figure 6, Movie S1)**. Calculation of the Gag lattice coverage from twenty-five HIV-2 immature particles revealed that the Gag coverage area underneath the viral membrane was as high as ∼90% (**Figure 5C, S5, Table S3)**, with the average membrane coverage ratio being 76 +/- 8%. The number of Gag molecules in the Gag lattice of these particles ranges from ∼1700 to ∼5700, with the average being ∼3900 +/- 1000 **(Figure 5D, Table S3**). The StA average of the Gag hexamer was carried out on ordered Gag hexamer positions by Dynamo [43, 44] and RELION-4.0 [45] **(Figure S3)**. The final reconstruction map **(Figure 5E, S4)** at 9.1 Å resolution revealed consistent structural features shown in the 5.5 Å map **(Figure 1)** that were computed by using the SPR method.

**Figure 6.**
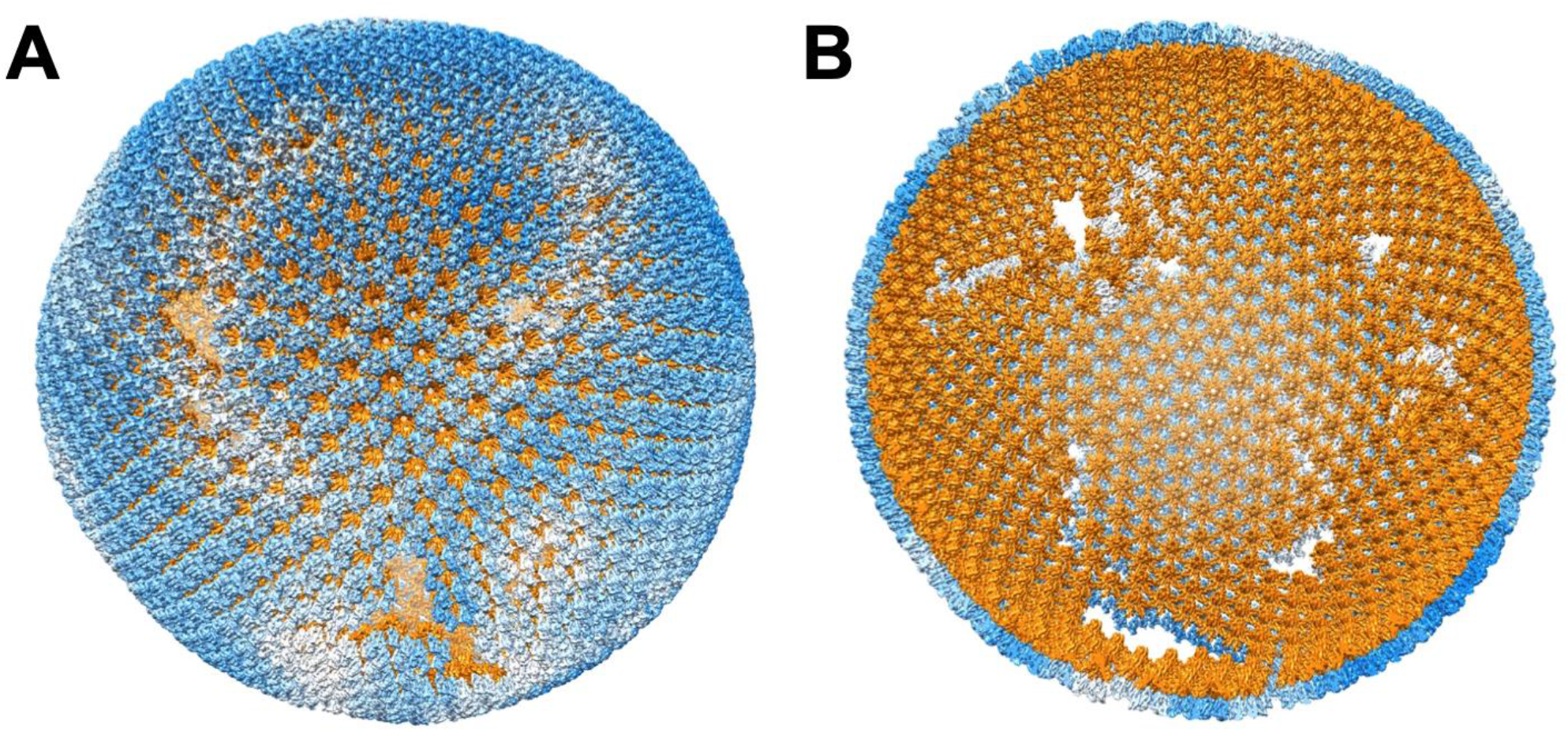
A model of HIV-2 immature Gag lattice structure. The model is generated by placing the 5.5 Å CA density onto the Gag lattice determined by Dynamo. (A) Exterior of the HIV-2 CA organization in the Gag lattice. Blue represents the NTD of the CA proteins. White indicates the interspersed gaps of the lattice. (B) Interior of the HIV-2 CA organization. Orange represents the CA_CTD_ and the CA-SP1 domains. The figure is produced using the UCSF Chimera [69] plug-in, Place Object [50].

To validate the observation of the Gag lattice was not due to the Gag expression system, we generated authentic immature HIV-2 particles (**Figure S6**). To do this, critical residues in the catalytic domain of the HIV-2 protease (PR) were mutated (*i.e.,* D25N/T26A/E37K) [46] in a HIV-2 ROD10 infectious molecular clone, and this was used to transiently transfect HEK293T/17 cells in order to generate authentic, non-infectious HIV-2 immature particles. Immunoblot analysis was performed on the released particles to confirm that no Gag processing had occurred, which was interpreted as indicating that the viral protease had been inactivated (**Figure S6A**). Cryo-EM analysis of these particles confirmed that authentic, immature HIV-2 particles could also possess an intact and near complete immature CA lattice (**Figure S6B**).

## Discussion

A previous study utilized a parallel comparative analysis to investigate immature particle morphologies among viruses representing the various retroviral genera [15]. Notably, it was observed that there were differences in the general subcellular distribution of the various retroviral Gag proteins as well as their being differences in the general morphology of the released immature particles. Furthermore, differences were observed among both the *Orthoretrovirinae* as well as the representative member of the *Spumaretrovirinae*. Intriguingly, among the observations made was that HIV-2 immature particles possessed a narrow range of particle size and had consistent regularly distributed electron density below the viral membrane, suggesting a tightly packed Gag lattice – in contrast to what was observed with HIV-1 immature particles. The use of cryo-ET and StA allowed for determination that the Gag lattice coverage in HIV-2 immature could be as high as ∼90% (**Figure 5C, 6)**, in contrast to HIV-1 in which Gag lattice covers about 60% of the viral membrane [47, 48]. The differences in HIV-1 and HIV-2 pathogenicity provide the impetus for gaining deeper insights into the biological differences of these closely related lentiviruses, particularly in gaining new insights into retroviral assembly.

We used cryo-EM SPR to solve the immature HIV-2 Gag hexamer structure at 5.5 Å resolution. Fitting of HIV-2 CA_NTD_ and CA_CTD_ structures into the resolved density map demonstrated a wineglass-like architecture of the central hexamer that is very reminiscent to that of HIV-1 [30], equine infectious anemia virus (EIAV) [49] and murine leukemia virus (MLV) [50]. The HIV-2 Gag hexamer was found to be stabilized by conserved residues in H2 at the three-fold inter-hexamer interface, by H9 in the CA_CTD_ at the two-fold inter-hexamer interface, and by a 6HB formed by the CA-SP1 at the six-fold intra-hexamer interface. We also observed a strong central pore density between the base of the CA_CTD_ and the 6HB that is consistent with the structural co-factor IP6. The observation of density in the central pore that likely represents IP6 is consistent with observations from previously determined retroviral structures [31, 49, 51]. The role of IP6 in CA-CA and Gag-Gag self-assembly was predicted by the pioneering studies conducted by Rein and coworkers [52]. For HIV-2, it is known to contain conserved lysine residues found within two rings of six lysine residues at the C-terminal end of CA_CTD_ (K157 and K226, or Gag residues K292 and K361, in HIV-2) [49, 51, 53].

Amino acid variation between the HIV-1 and HIV-2 CA proteins (*i.e.,* ∼70% sequence identity) imply that subtle differences in the structural configuration of their Gag hexamer and lattice organization can have important implications for virus structure and particle infectivity. For example, at the Gag lattice three-fold interface, we found that exchanging HIV-1 and HIV-2 residues significantly reduced virus particle production, and diminished virus particle infectivity. Mutations of a non-conserved residue in helix 2 of the CA proteins – *i.e*., HIV-2 CA G38M and HIV-1 CA M39G – reduced by 50% virus particle production as well as eliminating virus infectivity [54]. Additionally, the HIV-2 CA N127E mutant, which introduced the HIV-1 encoding residue into HIV-2 CA, severely reduced particle production. It should be further noted that HIV-2 is insensitive to the HIV-1 maturation inhibitor bevirimat (BVM) [55, 56], emphasizing that subtle differences in structural property between the HIV-1 and HIV-2 Gag proteins can have important implications in retrovirus replication. The binding site of BVM has been determined to be inside the CA-SP1 six-helix bundle near the IP6 binding site [30, 57, 58]. Taken together, these observations argue for the importance in obtaining a near-atomic structure of HIV-2 immature Gag lattice to gain further mechanistic insights into the comparative differences between HIV-1 and HIV-2 CA structure.

From the HIV-2 CA_CTD_ crystal structure, it can be inferred that residues in H10 and H12 play a critical role in mediating the dimeric interface in the crystal. In particular, the C-terminal residues of the HIV-2 CA form a 3_10_ helix, *i.e.,* H12. The angle in which the HIV-2 CA_CTD_ H12 extends from H11 is different from that in our immature HIV-2 CA lattice from particles (**Figure S7 A-B**), as well as immature HIV-1 CA-SP1 lattice from in vitro assemblies or virus particles [30]. However, modeling of the HIV-2 CA_CTD_ crystal structure into the HIV-1 mature CA capsid implies that the putative H12 is in the vicinity of H10 in the adjacent hexamer at the three-fold CA inter-hexamer interface (**Figure S7 C-D**). Therefore, additional studies were conducted to assess the biological relevance of the residues in these helices.

Site-directed mutagenesis of the HIV-2 CA L199 (in H10), along with R228, and L229 (in H12) led to severe impairments in particle production and infectivity (**Figure 4 A-C**). This implies that the residues at the H10 and H12 revealed in the HIV-2 CA_CTD_ crystal may be important for HIV-2 morphology and particle infectivity, and help explain the morphological features distinct from that of HIV-1.

Cryo-EM images (**Figure 4D, S8**) show that mutations at a H10 residue L199 (L199K and L199G) did not significantly change particle production in the immature particles and immature Gag lattice features were prevalent within the particles.. No capsid cores were observed in the mature particles for L199K mutant. For L199G, less than two percent of observed particles have internal densities that are attributable to presence of a full or partial capsid core. However, the mutant is not infectious suggesting these cores are defective. Mutation at the H12 residue R228 (*i.e.,* R228E) did not eliminate Gag assembly. But the Gag densities in R228E immature particles appear different from normal Gag lattice in the WT HIV-1 and HIV-2 immature particles. Intriguingly, these lattice-like densities resemble the morphology of HIV-1-CA_NTD_/HTLV-1-CA_CTD_ chimera particles that we previously reported using similar analyses [59]. No capsid cores were observed in mature R228E particles. Presumably, because the HIV-2 CA residues E228 and L229 are part of the 6HB and are 3 and 2 amino acids upstream of the CA-SP cleavage site (*i.e.,* CA M230 and SP1 A1, respectively), it is possible that mutations at these residues impair proper Gag assembly and protease cleavage, thus preventing the formation of the mature CA lattice. Nevertheless, the phenotypes of mutations on the H10 residue L199 are clearly novel and not postulated by the current lentivirus CA structural model. For example, the aligned residue of HIV-2 L199 in HIV-1 H10 (*i.e.,* T200) is not directly involved in any molecular interaction in either immature Gag (*e.g.,* PDB 5L93) or mature CA lattice structures (*e.g*., 5MCX and 6SKN). Determination of the high-resolution structure of mature HIV-2 CA lattice and understanding the implications of the mutagenesis studies on HIV-2 replication in mechanistic basis of HIV-2 maturation represent an important area for future studies.

## Materials and Methods

### HIV-2 CA_CTD_ protein purification, crystallization, and structure determination

We expressed HIV-2 CA_CTD_ (residues 145-230) in *E. coli* strain BL21 (DE3) with a N-terminal 6xHis-tag, using the pET28a expression vector. The expressed protein was purified by Ni-NTA affinity and Superdex 75 size-exclusion chromatography (SEC) steps. The 6xHis-tag was removed by a treatment with HRV 3C protease after elution from the Ni-NTA column and prior to the SEC step. The purified protein in 20 mM Tris-HCl (pH 7.4), 0.5 M NaCl, + 5 mM β-mercaptoethanol was concentrated by ultrafiltration to ∼43 mg/ml. Crystallization screening was performed on a Phoenix (Art Robbins Instruments, USA) crystallization machine. Crystals appeared in one day and grew to full size in three days. The crystals that gave the best diffraction were grown under the condition of 1.2 M tri-sodium citrate, 0.1 M Bis-Tris propane, pH 7.0.

The HIV-2 CA_CTD_ crystals were cryo-protected by soaking in the crystallization reservoir solution supplemented with 25% ethylene glycol and flash cooled in liquid nitrogen. X-ray diffraction data were collected at the 24-ID-E (or C) beam line of the Advanced Photon Source, using the wavelength of 0.979 Å and processed using HKL2000. The crystal structure was solved by the molecular replacement phasing, using the coordinates of 6AXX (PDB ID: 6AXX) as the search model. Iterative model building and refinement were performed using COOT [60] and PHENIX [32] giving a final Rwork of 22.39% and Rfree of 24.95%, respectively. The final model consists of 84 amino acids. Refinement and validation metrics are listed in **Table S1**.

### Immature particle production and purification

The HIV-2 *gag* gene (bases 546 to 2114 from GenBank accession number M15390) was used as the basis to create a codon-optimized *gag* gene that was cloned into the pN3 expression vector that included within a cassette that contains a 5′ Kozak sequence (GCACC**ATG**, start codon in bold). The pN3-gag expression constructs and their homologous envelope glycoprotein (gp160) expression plasmids were co-transfected into HEK293T/17 or 293F cells using GenJet Ver. II (SignaGen Laboratories, MD) at a mass ratio of 10:1. After 48 h (for HEK293T/17 cells) or 120 h (for 293F cells) post-transfection, the cell culture supernatant was harvested and filtered through a 0.22 μm filter. Purification of the particles was conducted by a series of ultracentrifuge steps. Initially, the viral supernatant was pelleted over an 8% (v/v) OptiPrep^TM^ cushion in STE buffer (100 mM NaCl, 10 mM Tris-Cl pH 7.4, 1 mM EDTA) with a 50.2 Ti rotor (Beckman Coulter, CA) at 35,000 rpm for 90 min. The pellet was resuspended in STE buffer and banded out over an OptiPrep^TM^ step gradient (10, 20, 30% OptiPrep^TM^ (v/v)) with a SW55 Ti rotor (Beckman Coulter, CA) at 45,000 rpm for 3 h. The final particle sample was suspended in STE buffer.

### Mature particle production

The HIV-2 *pROD-MIG* plasmid and a VSV-G expression construct were co-transfected into HEK293T/17 cells using GenJet Ver. II (SignaGen Laboratories, MD) at a 3:1 mass ratio to produce WT and mutant infectious viral particles. After 48-hours post-transfection, the viral supernatants were harvested, clarified by centrifugation (1,800 × g for 10 min), and filtered through 0.22 µm filters. Then the supernatants were concentrated by ultracentrifugation in a 50.2 Ti rotor (Beckman Coulter, CA) at 35,000 rpm for 90 min through an 8% OptiPrep^TM^ (Sigma-Aldrich, MO) cushion. The efficiency of mature particle production was analyzed by quantifying the CA (p24) band in culture supernatants using anti-HIV-1 p24 primary antibodies (NIH HIV Reagent Program, Manassas, VA – Mouse anti p24: ARP-6521) and goat anti mouse IRDye® 800CW (LI-COR Biosciences, NE). Capsid protein levels were assessed by SDS-PAGE followed by immunoblotting with an anti-HIV-1 CA (p24) antibody. CA expression levels were normalized relative to GAPDH levels, and the mutant Gag expression levels were determined relative to that of WT HIV-2 Gag.

### Authentic HIV-2 particle production

The HIV-2 ROD10 infectious molecular clone was modified by introducing several mutations in the HIV-2 protease active site (*i.e.,* D25N/T26A/E37K) [46]. Particles were produced in HEK293T/17 cells in the same fashion as the mature particles. Cell lysates and viral supernatants were collected and immunoblotted for HIV-2 Gag using a p24 antibody to confirm a full-length intact Gag (**Figure S6**).

### Quantification of Gag and CA proteins in WT and mutant viruses

The Gag or CA protein concentrations of the immature or mature HIV-2 particles were first measured by using the BCA assay (Pierce, Rockford, IL). The samples were subjected to SDS-PAGE and then transferred to nitrocellulose membranes. The Gag and CA proteins from immature or mature particles were detected with a 1:1,500 dilution of mouse anti-HIV p24 antibody (Catalogue #: ARP-6521; NIH HIV Reagent Program, Manassas, VA) in 5% milk TBST (Tris-buffered saline plus Tween- 20).Membranes were washed before incubation with a goat anti-mouse 1:5000 IRDye® 800CW secondary. To compare the Gag or CA protein quantification of WT to that of mutant, 293T/17 cells were collected and lysed with Cellytic™ M cell lysis buffer buffer (Sigma-Aldrich, MO) and clarified via centrifugation (1800 x g for 10 min). The Gag and CA proteins in cell lysates were detected with a 1:2,000 dilution of mouse anti-HIV p24 antibody (Catalogue #: ARP-6521; NIH HIV Reagent Program, Manassas, VA) and 1:10,000 rabbit anti-GAPDH antibody (Catalogue #: ab128915; Abcam, Cambridge, United Kingdom) in 5% milk TBST. Membranes were washed before incubation with a 1:5000 dilution of goat anti-mouse IRDye® 800CW and a 1:5000 dilution of goat anti-rabbit IRDye® 680RD secondary. The membranes of the immunoblots were imaged by using a ChemiDoc Touch system (Bio-Rad, CA). The densities of the protein bands were measured by using ImageJ. To access particle production (i.e., particle release) of mutants relative to that of WT, the Gag or CA expression levels detected from cell lysates of the WT and mutants were normalized relative to the respective GAPDH level. Mutant particle release to that of WT was determined by the ratio of Gag levels detected from cell culture supernatants to that from the normalized Gag levels from the cell lysate, with WT particle release being set to 100 and particle release of the mutants being relative to that of the WT. Results were plotted by using GraphPad Prism 6.0 (GraphPad Software, Inc., CA). Relative significance between a mutant and WT was determined by using an unpaired t-test. Immunoblot analyses were done in three independent replicates to access either immature particle production (**Figure 4A, Figure S9**) or mature particle production (**Figure 4B, Figure S10**). To confirm the immunoblot analysis done with anti-CA antibodies was not influenced by the CA mutations, control experiments were done using an anti-HIV-1 p17 antibody (Cat. No. sc-69723, Santa Cruz Biotech, TX) (**Figure S11**).

### Infectivity assay

The HIV-2 *pROD-MIG* plasmids and the VSV-G expression construct were co-transfected into HEK293T/17 cells using GenJet, ver II (SignaGen, Gaithersburg, MD) at a 3:1 ratio to produce WT and mutant infectious viral particles. After 48-hours post-transfection, the viral supernatants were harvested, clarified by centrifugation (1,800 × g for 10 min), and filtered through 0.22 µm filters. U373-MAGI- CXCR4 cells were plated in a 12-well plate and each was treated with 1 ml viral supernatants and 1 mL fresh medium. Each group had 4 well replicates. The cells were collected for fluorescence analysis via BD LSR II flow cytometer (BD Biosciences, NJ) 48-hours post-infection as described before [53]. Flow cytometry data were examined in FlowJo v.7 (Ashland, OR). The infectious cells were calculated from the flow data by adding all positive quadrants (mCherry+ only, GFP+ only, and mCherry+/GFP+) to determine infectivity. Mutant infection level was determined related to WT. Then, the relative infectivity of each group was normalized to its relative mature particle production as assessed by p24 immunoblot of the produced particles. Three independent experiments were performed (**Figure 4C**).

### Transmission electron microscopy (TEM) sample preparation

The frozen-hydrated TEM grids are prepared by manual blotting in a FEI MARK III Vitrobot system. A small volume (∼3.5 μl) of concentrated particle sample, or particle mixed with BSA-treated 20nm Nano gold particles, were applied to a glow-discharged Quantifoil holey carbon grid (Ted Pella, Redding, CA) and then blotted with a filter paper before being plunged into liquid ethane. The frozen grids were first examined and screened in an in-house FEI TF30 field emission gun transmission electron microscope at liquid nitrogen temperature (FEI Company, Hillsboro, OR).

### Single particle data collection and reconstruction

The cryo-EM data used for image processing were collected on a Titan Krios TEM at the National Cryo-EM Facility, Cancer Research Technology Program, Frederick National Laboratory for Cancer Research. The images were recorded using a K2 Summit direct electron detector that was positioned post a Gatan BioQuantum energy filter with a 20 eV energy slit for the zero-loss image collection. The cryo-EM movies were recorded at 105,000x nominal magnification, which corresponds to 0.66 Å at the specimen in super resolution mode. Each movie contains 40 frames with an accumulated electron dose of 30-50 e^-^/Å^2^. The defocus levels for the cryo-EM data collection were set to 0.75 to 2.25 μm. The whole data set contains 2508 movies. Movies were motion corrected and dose-weighted using the MotionCor2 program [61]. The contrast transfer function (CTF) parameters were determined using Gctf [62]. Detailed data acquisition parameters are listed in **Table S2**.

The single particle reconstruction was processed in RELION-3.1 **(Figure S1A)** [63, 64]. Initially, ∼1,000 Gag lattice regions on the edges of the particles were picked out from 2415 micrographs to generate initial 2D averages as templates. Then, 219,321 particles were identified by using a template-based auto picking procedure implemented in RELION-3.1. The box size of these local Gag lattice regions was 400 pixels, corresponding to 528 Å on the specimen. The extracted particles were subjected to multiple cycles of 2D classification (remove contaminates and disordered classes) and 3D classification (identify a homogenous set of particles). The final reconstruction from a selected 46,017 particles.

To conduct the Fourier-shell correlation coefficient (FSC) analysis, a customized routine was developed to sort the particles into two non-overlapping data sets of the roughly equal number of particles according to their locations in the micrographs **(Figure S1B)**. Briefly, the particles within each micrograph were first separated into multiple clusters, with the distance between any two clusters larger than a given threshold. Next, clusters in all micrographs were assigned to one of the two data sets alternatively based on the cluster size, to maintain roughly equal number of particles between the two data sets. The algorithm was implemented in the MATLAB programming language. This method avoids assigning the overlapped regions of the crystalline Gag lattices into the same half data set. The final resolution was determined to be 5.5 Å at the FSC cut-off of 0.143 **(Figure S2**).

### Gag lattice model building and refinement

The HIV-2 CA-SP1 protein structural model is generated from three components. The NTD (Gag residues:S150 to Y279) from HIV-2 NTD structure (PDB ID: 2WLV), CTD (T282 to C352, the crystal structure), and CA-SP1 (Q353 to I373) from threading the HIV-2 ROD CA sequence over the immature HIV-1 Gag cryo-EM structure (PDB ID: 5L93) [30]. The residues N280 and P281 were added manually to complete the sequence. The protein models were generated using by performing iterative cycles of the real-space refinement routines in Phenix [32] and manual perturbations in Coot [60].

### Cryo-ET reconstruction and subtomogram averaging

Cryo-ET reconstruction of immature HIV-2 particles, HIV-2 MIG construct, and its mutant was carried out by using IMOD [65] and EMAN2 [66] respectively. The microscope defocus levels for the tilt series were determined by CTFFIND4 [67]. The CTF corrected reconstruction maps of immature HIV-2 particles were used for further StA analysis. The StA of HIV-2 Gag hexamers of 20 immature particles from five tilt series was carried out by an approach using Dynamo [68] and RELION-4.0 [45] (**Figure S3**).

The StA procedure using Dynamo has been previously described in detail [43] . Briefly, the annotated centers and radii of the particles [68] were used to generate an initial sampling of 12,109 points of all particles. These sampling points are regularly distributed on a sphere of the measured radius of each particle with an average distance of 92 Å between closest positions [65]. Subtomograms with a box size of 256 pixels (338 Å) were cropped on the sampling positions and were assigned with initial orientations along the normal direction to the surface of the particle with randomized azimuth angles. The averaged map from the particles within one tilt series was used as the starting model for the initial alignment. The relative orientations and positions of the subtomograms from all tilt series were then calculated using C1 symmetry. The resulting map was then shifted so that the center of the Gag hexamer was aligned with the z axis of the model. The new map was then used as the 2^nd^ model for further refinement with C6 symmetry. Next, for each particle, a new set of 19 sampling points are generated according to the relative positions of the Gag hexamers in the lattice. The subtomogram data set was expanded to 37,081 subtomograms for the total 20 immature HIV-2 particles. The new set of subtomograms with a box size of 192 pixels (253 Å) were extracted from the IMOD reconstruction maps. Further refinement resulted in a converged map and Gag lattice positions computed from 13,378 subtomograms.

A two-step selection procedure was employed to further narrow down the subtomograms prior to refinement by using RELION-4.0. First, a custom MATLAB algorithm was developed to clean up the small amount of data points that were in the membrane envelope. An estimated smooth and robust Gag lattice surface was first generated by multiple cycles of computation that fits each point and its surrounding points to a spherical cone and rejects the outlier point. The statistical distribution of the distance between each data point to the theoretical surface was then used to determine a threshold value that filtered out the spurious outer shell points. A set of particle data points (14,675 points) with the orientation within 15 degrees from the norm of the modeled Gag lattice surface were selected. Next, the UCSF Chimera [69] plug-in routine, Place Object [50], was used to display the Gag lattice points color-coded by the correlation coefficient (CC) values between the sub-volumes with the averaged map **(Figure 5B)**. A CC threshold was determined by eye for each tomogram to remove the data points on the edges of the continuous Gag lattice patch resulting in the data set of the ordered lattice regions. This procedure resulted into 10,333 positions that were subjected to further refinement in RELION-4.0 [45]. Pseudo-subtomograms with a box size of 256 pixels (338 Å) were extracted from the original tilt series. A *de novo* model was generated and used for calculating the orientations and positions to produce an average map with C6 symmetry. The tomo CTF and frame alignment were carried out in RELION-4.0 and the final reconstruction map was determined to be 9.1 Å based on the Fourier Shell correlation cut-off of 0.143 (**Figure S4)**.

### Coverage of Gag molecules underneath the viral membrane

The surface area covered by Gag molecules organized in hexagonal lattice was estimated for each HIV-2 immature virus particle. The dense seed locations of 10 pixels distance (13.2 Å) on the Gag lattice layer were generated. Subtomograms with a box size of 100 pixels (132 Å) cropped from the tomograms at these positions contain at least one Gag hexamer within the box. The resulting 624,669 boxes (from 25 immature HIV-2 particles) were realigned using Dynamo [44, 68] allowing them to shift freely around the surfaces of the particles. After alignment, duplicates of final positions representing the same physical particle were located by DBSCAN [63] and removed, producing a clean data set of unique box positions **(Figure S5A)**. The distribution pattern of the candidate Gag hexamer centers around each data point was investigated through the “neighborhood analysis” method [37]. In this approach, the relative distances and angles were computed for every pair of aligned subtomograms present in the data set, showing that the computed positions for the neighbors of a given Gag hexamer are typically distributed in a local hexagonal lattice with a step of 50 pixels (66 Å). After observing the emergence of a hexagonal pattern, the subtomograms that had at least 4 closest neighbors at the distance imparted by this pattern were kept **(Figure S5B-C)**. The total number of Gag molecules involved in formation of the Gag lattice in the immature virus particle is estimated to be six times of the total number of remaining subtomograms, which is also used to compute the percentage of the particle surface covered by the Gag hexamers **(Figure S5D, Table S3)**.

### Data availability

The refined model of HIV-2 CA_CTD_ crystal structure was deposited in the protein data bank (PDB ID: 7TV2). The 5.5 Å cryo-EM SPR map of the HIV-2 Gag lattice, the 9.1 Å cryo-ET/StA map, and the hexamer model fitted into the 5.5 Å SPR map have been deposited in the EMDB and PDB with accession codes EMD-29607, EMD-29606, and PDB ID: 8FZC respectively.

## Acknowledgements

This work is supported by NIH grant R01 AI177264 (to L.M. and J.M.). Support is also acknowledged from NIH grants R35 GM118047 (to H.A.) and R21 AI148328 (to W.Z. and L.M). D.C.D. acknowledges the support of the grant 205321/179041 of the Swiss National Science Foundation (SNF), the grant RGP0017/2020 of the Human Frontiers Science Program (HFSP), and funding from the project PID2021-127309NB-I00 funded by AEI/10.13039/501100011033/ FEDER, UE. We thank the staff at the National Cryo-EM Facility, Cancer Research Technology Program, Frederick National Laboratory for Cancer Research for high resolution cryo-EM and cryo-ET data collection. Parts of this work were carried out in the Characterization Facility, University of Minnesota, which receives partial support from the NSF through the MRSEC (Award Number DMR-2011401) and the NNCI (Award Number ECCS-2025124) programs. Cryo-ET data collection of HIV-2 MIG construct, and mutants were carried out by Dr. Xiaofeng Fu at the Biological Science Imaging Resource at Florida State University through the U24 GM116788 program. X-ray diffraction data were collected at the Northeastern Collaborative Access Team beamlines, which are funded by the US National Institutes of Health (NIGMS P30 GM124165). Cryo-ET StA computations were carried out on the GPU clusters at Minnesota Supercomputing Institute. N.T. was supported by NIH grants T32 DA007097, F32 AI150351, and American Cancer Society Postdoctoral Fellowship PF-21-189-01-MPC. W.G.A. was supported by the Institute for Molecular Virology Training Program (i.e., NIH grant T32 AI83196).

## Author contributions

L.M.M. and W.Z. designed the research. N.T. and L.M. Mendonça purified the immature HIV-2 particles. N.T. performed cryo-EM and cryo-ET reconstructions. K.S. and H.A. solved the crystal structure of HIV-2 CA_CTD_. H.Y., S.M. and W.G.A., performed mutagenesis studies. H.Y. and G.C.B. analyzed production and performed infectivity assays of HIV-2 WT and mutant viruses. H.Y. and S.M. produced mutant viruses for cryo-EM examination. N.T. and W.Z. imaged the WT and mutant virus samples by cryo-EM. W.Z., R.C. and D.C.D. analyzed the Gag lattice structure by the cryo-ET StA method. R.C. and D.C.D. computed the surface coverage of Gag lattices using the DBSCAN and neighborhood analysis methods. G.Y. wrote several imaging processing and analysis programs that enhanced quality and efficiency of the computation. N.T. and K.S. performed structural modeling and fitting. All authors contributed to either drafting and/or revising the paper.

---

**Table S1.** Data collection and refinement statistics.

|  | HIV-2<sub>CTD</sub> |
| --- | --- |
| **Data Collection** |  |
| Wavelength (Å) | 0.979 |
| Resolution range (Å) | 24.4 - 1.98 (2.05 - 1.98) |
| Space group | C2 |
| Unit cell |  |
| *a,b,c* (Å) | 48.813, 30.919, 46.494 |
| $\alpha,\beta,\gamma$ (°) | 90, 90.561, 90 |
| Total reflections | 13142 (1335) |
| Unique reflections | 4779 (489) |
| Multiplicity | 2.7 (2.7) |
| Completeness (%) | 95.44 (95.66) |
| *I*/$\sigma$(*I*) | 11.41 (3.29) |
| R-merge | 0.0624 (0.267) |
| R-meas | 0.0762 (0.326) |
| R-pim | 0.0430 (0.185) |
| CC<sub>1/2</sub> | 0.997 (0.963) |
| **Refinement** |  |
| Reflections used | 4761 (485) |
| # for R-free | 237 (27) |
| R-work | 0.2239 (0.3252) |
| R-free | 0.2495 (0.3944) |
| Number of non-hydrogen atoms | 659 |
| Macromolecules | 644 |
| Solvent | 15 |
| Protein residues | 83 |
| RMSD |  |
| Bond lengths (Å) | 0.003 |
| Bond angles (°) | 0.60 |
| Ramachandran plot |  |
| Favored (%) | 97.53 |
| Allowed (%) | 2.47 |
| Outliers (%) | 0.00 |
| Average B-factor | 49.45 |
| Macromolecules | 49.62 |
| Solvent | 42.34 |

Statistics for the highest-resolution shell are shown in parentheses.

**Table S2.** Cryo-EM data collection.

| **Imaging Mode** | **Single Particle** | **Cryo ET** |
| --- | --- | --- |
| ***Data Collection*** |  |  |
| Microscope | FEI Titan Krios | FEI Titan Krios |
| Voltage (kV) | 300 | 300 |
| Magnification | 105,000× | 105,000× |
| Energy-filter (eV) | 20 | 20 |
| Detector | Gatan K2 Summit | Gatan K2 Summit |
| Recording Mode | Super-resolution | Super-resolution |
| Pixel size (Å/pixel) | 1.32 | 1.32 |
| Defocus range setting (μm) | 0.75 to 2.25 | 2.5 |
| Total Dose (e<sup>-</sup>/Å<sup>2</sup>) | 50-60 | 50 |
| Dose rate (e<sup>-</sup>/sec/pixel) | 8.7 | 1.1 |
| Exposure time (sec) | 10.0 | 2.0 |
| Frame number | 40 | 8 per tilt |
| Dataset size | 2508 micrographs | 63 tilt series |
| Tilt range | N/A | ±60 |
| Tilt step | N/A | 3 |

**Table S3.** HIV-2 lattice morphology.

| **Index** | **Tomogram** | **N. of initial sampling points** | **Radius (nm)** | **Surface area coverage (%)** | **N. of Gag molecules** |
| --- | --- | --- | --- | --- | --- |
| 1 | HIV2ts3_02 | 31067 | 66 | 88 | 5364 |
| 2 | HIV2ts3_02 | 30220 | 64 | 78 | 4734 |
| 3 | HIV2ts3_02 | 36533 | 71 | 78 | 5724 |
| 4 | HIV2ts3_02 | 14930 | 45 | 63 | 2100 |
| 5 | HIV2ts3_02 | 27651 | 62 | 70 | 3996 |
| 6 | HIV2ts3_02 | 17142 | 62 | N/A | 2670 |
| 7 | HIV2ts3_03 | 36909 | 71 | 76 | 5532 |
| 8 | HIV2ts3_03 | 33605 | 68 | 75 | 5112 |
| 9 | HIV2ts3_03 | 22931 | 56 | 72 | 3378 |
| 10 | HIV2ts3_03 | 23501 | 57 | 83 | 3702 |
| 11 | HIV2ts3_03 | 23053 | 56 | 66 | 3378 |
| 12 | HIV2ts3_27 | 26151 | 60 | 79 | 4104 |
| 13 | HIV2ts3_27 | 24491 | 58 | 74 | 3666 |
| 14 | HIV2ts3_27 | 27088 | 61 | 83 | 4284 |
| 15 | HIV2ts3_27 | 21712 | 55 | 89 | 3624 |
| 16 | HIV2ts3_27 | 22541 | 56 | 76 | 3468 |
| 17 | HIV2ts3_27 | 32274 | 69 | N/A | 4956 |
| 18 | HIV2ts3_37 | 23008 | 56 | 78 | 3702 |
| 19 | HIV2ts3_37 | 26910 | 61 | 80 | 4302 |
| 20 | HIV2ts3_37 | 21962 | 55 | 75 | 3384 |
| 21 | HIV2ts3_37 | 12483 | 42 | 61 | 1674 |
| 22 | HIV2ts3_49 | 25778 | 60 | 88 | 4398 |
| 23 | HIV2ts3_49 | 18761 | 51 | 59 | 2466 |
| 24 | HIV2ts3_49 | 20685 | 61 | N/A | 3720 |
| 25 | HIV2ts3_49 | 23272 | 57 | 81 | 3768 |

Note:
- 1) Particles #6, 17, and 24 are not full particles in the tomograms and were excluded from the calculation of the surface area coverage. - 2) The radius of the Gag lattice is defined as the radius of the linker region (between CA<sub>NTD</sub> and CA<sub>CTD</sub>) of the CA density layer.

**Movie S1.**
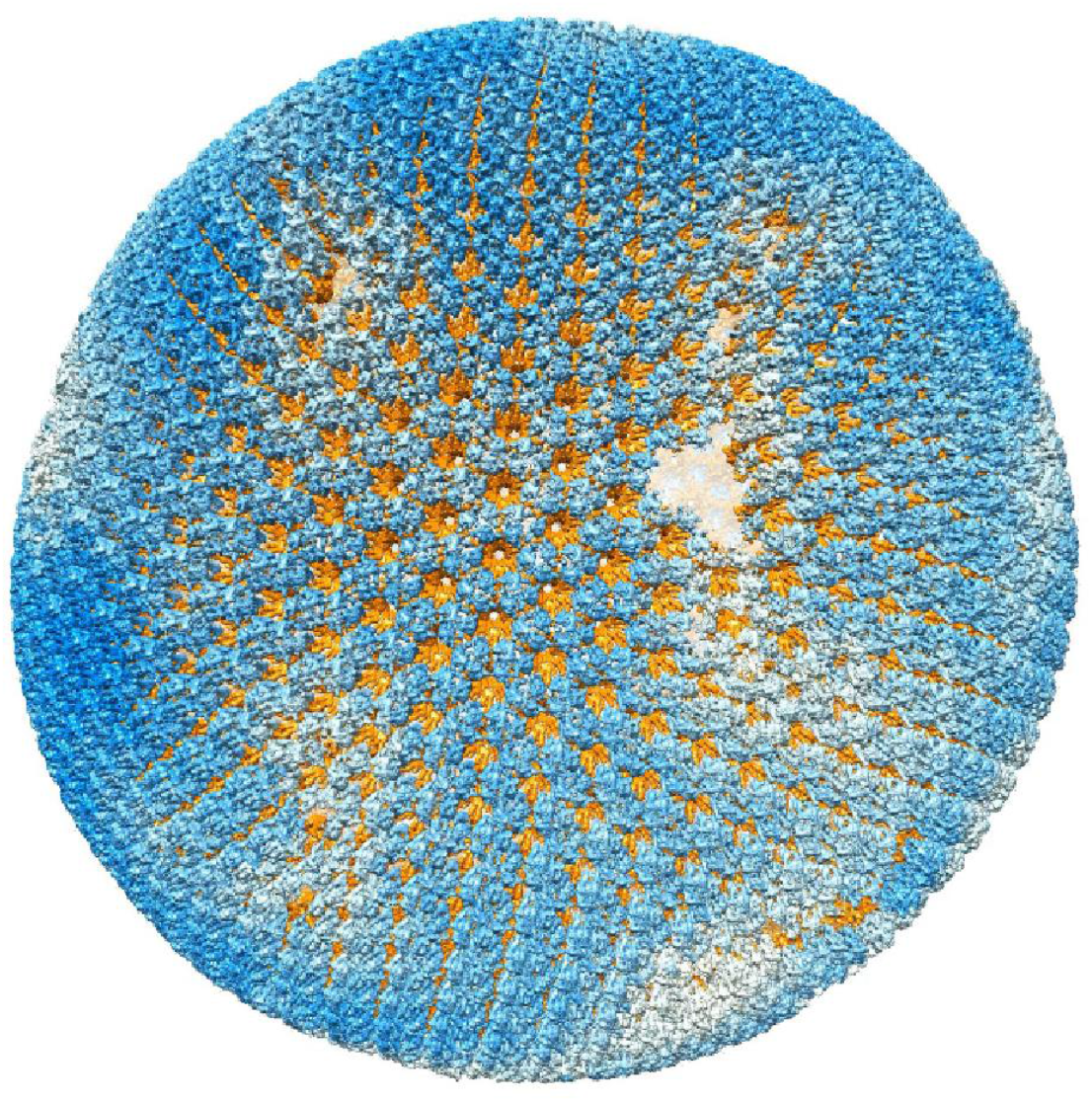
Model of a HIV-2 immature particle. The model is generated by placing the 5.5 Å CA hexamer density onto the Gag lattice determined by the StA Dynamo Gag hexamer position calculations. The NTD of the CA proteins are colored in blue and white. The CTDs and the CA-SP1 domains are colored in orange. The color denotes how well that hexamer matches the StA model, with blue having a higher correlation coefficient score than white.

**Figure S1.**
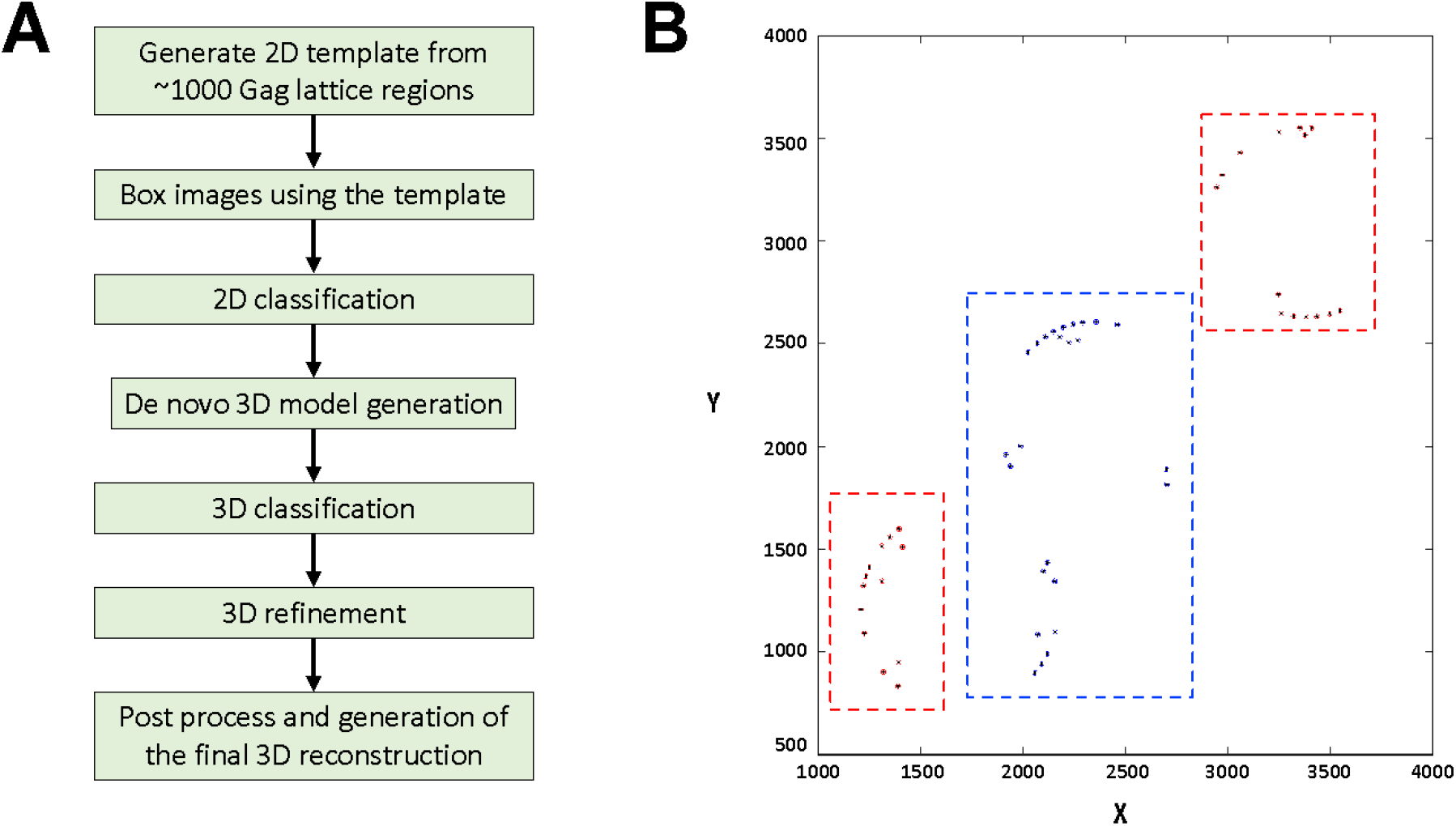
Workflow of cryo-EM single particle reconstruction. (A) The steps involved in calculating the 5.5 Å reconstruction map of HIV-2 Gag lattice structure. (B) Illustration showing the coordinates of the boxed particles sorted into two groups (blue and red) according to their locations in the micrograph. The two half sets are used for for resolution determination according to gold standard refinement procedures [70]

**Figure S2.**
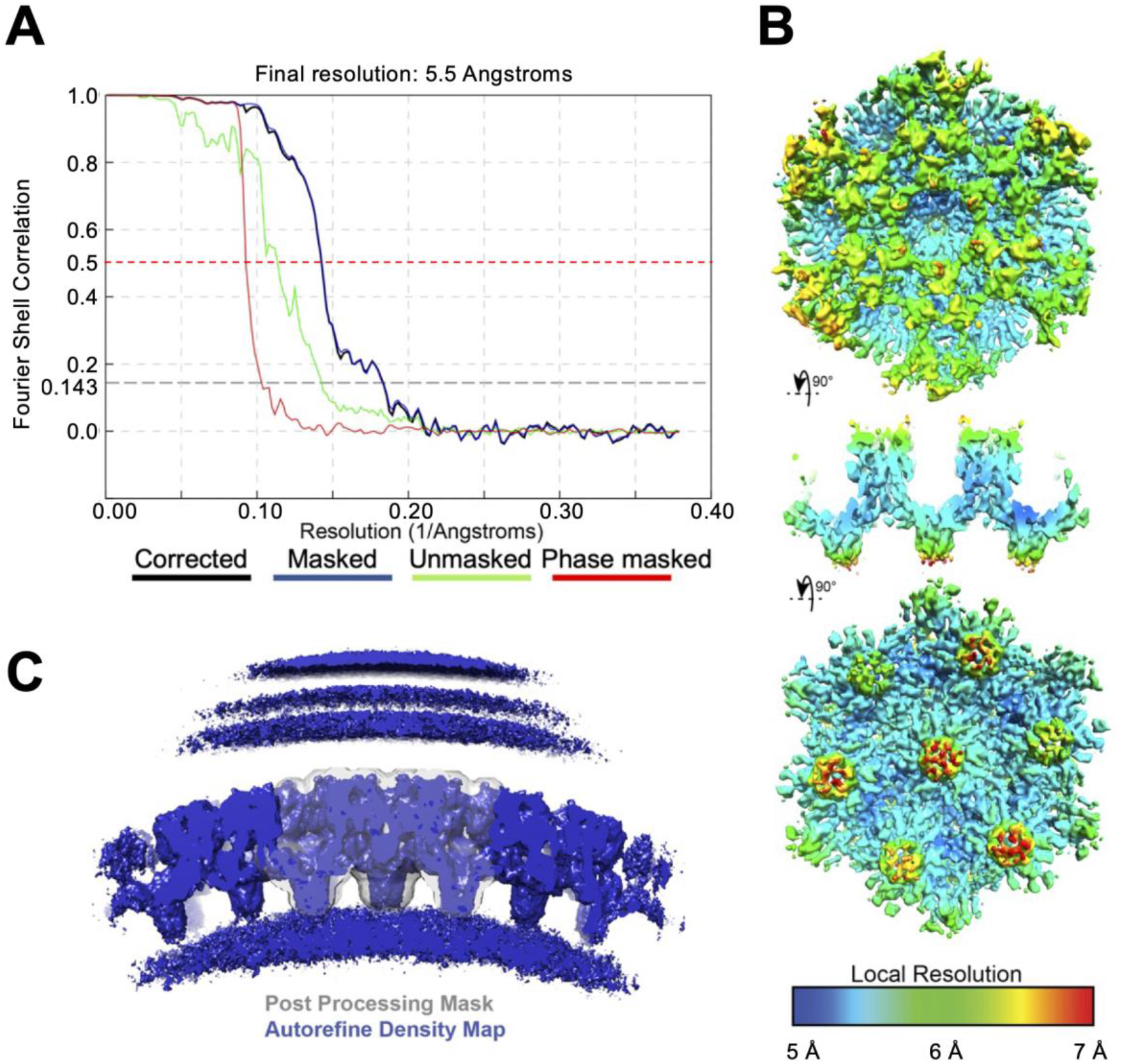
Cryo-EM structure validation of SPR of HIV-2 immature particles. (A) Gold standard FSC curves. (B) LocalRes result for the CA density in the reconstruction. (C) Central slice through the unmasked reconstruction density from 3D Autorefine along with CA mask used for generating the FSC curve in panel A.

**Figure S3.**
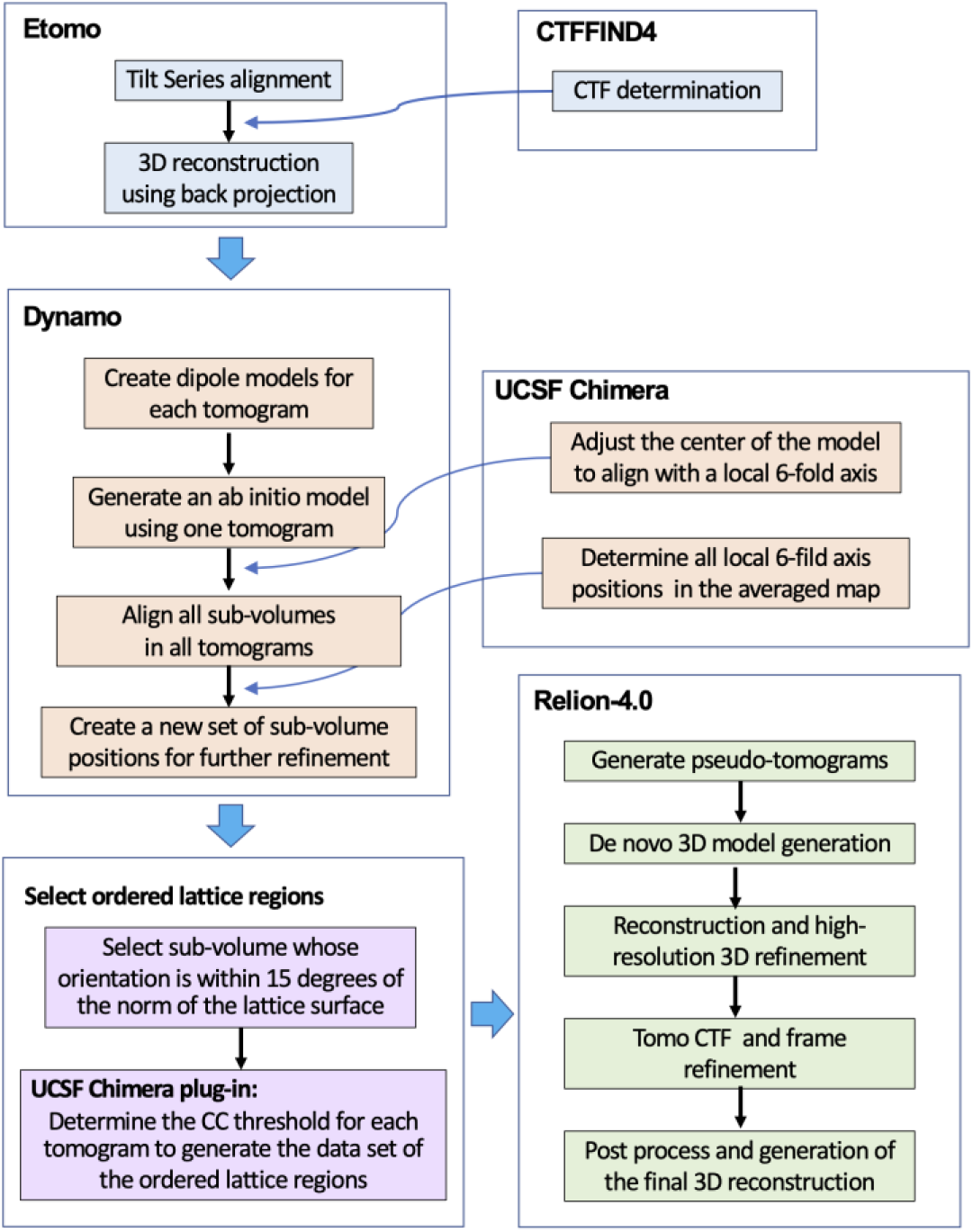
Workflow of cryo-ET and sub-tomogram averaging procedure. The software packages used in the cryo-ET and sub-tomogram averaging procedures include IMOD (Etomo) [65], CTFFIND4 [67], Dynamo [43], and RELION-4.0 [45]. The major computation steps for each software package are outlined in the respective box. The thick blue arrows represent the workflow of the sequential computational steps. The thin blue arrows represent information from CTFFIND4, and UCSF Chimera [69] is respectively integrated in the Etomo and Dynamo computations. The UCSF Chimera plug-in routine, Place Object [50], was used to determine the CC threshold for each tomogram to generate the data set of the ordered lattice regions.

**Figure S4.**
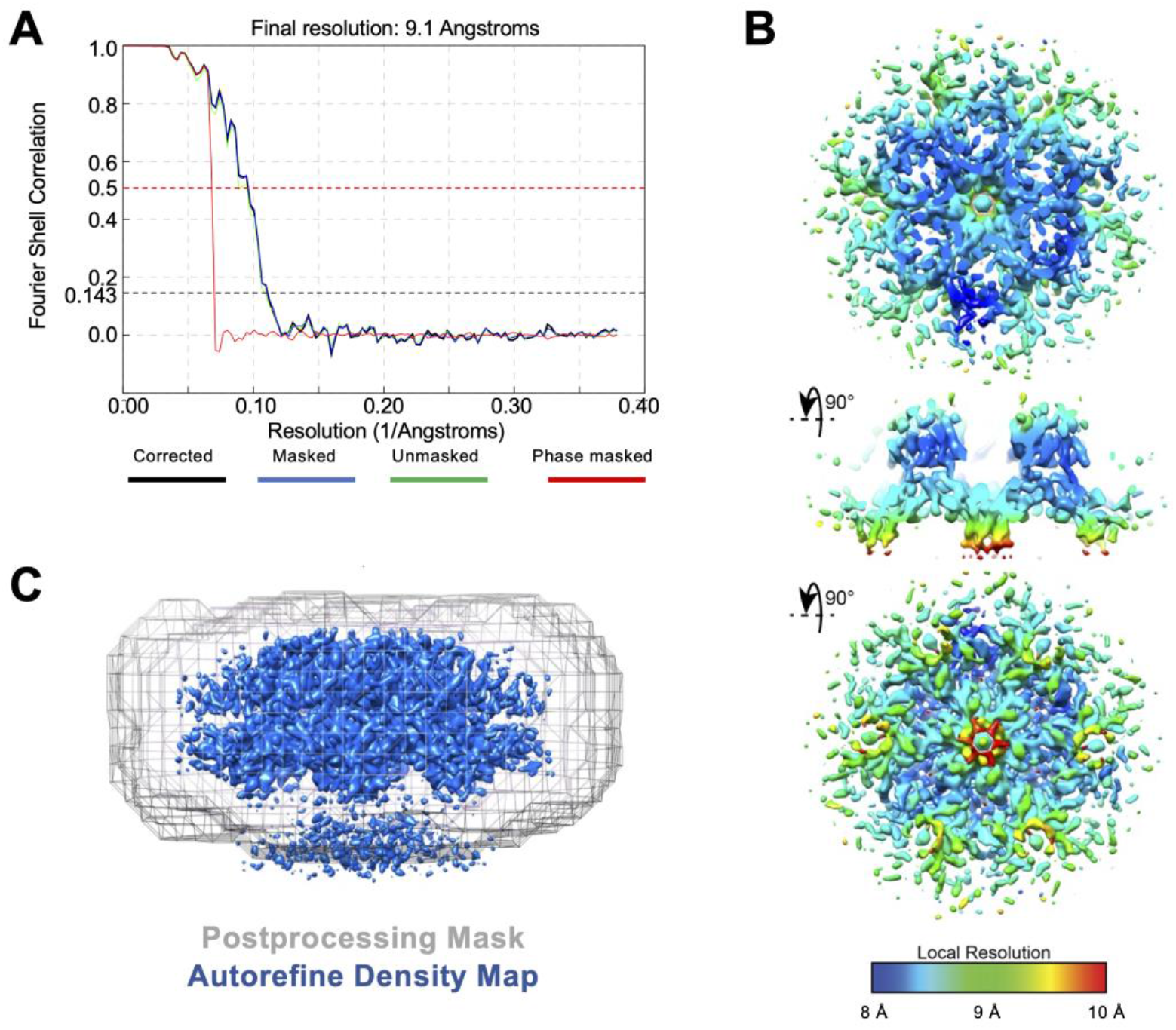
Resolution validation of HIV-2 immature particle Gag hexamer structure determined by the cryo-ET and sub-tomogram averaging methods. (A) Gold standard FSC curves. The resolution was determined to be 9.1 Å using FSC cut-off to be 0.143. (B) LocalRes result for the CA density in the reconstruction. (C) Sideview of the unmasked reconstruction density from 3D Autorefine along with the mask used for generating the FSC curve in panel A.

**Figure S5.**
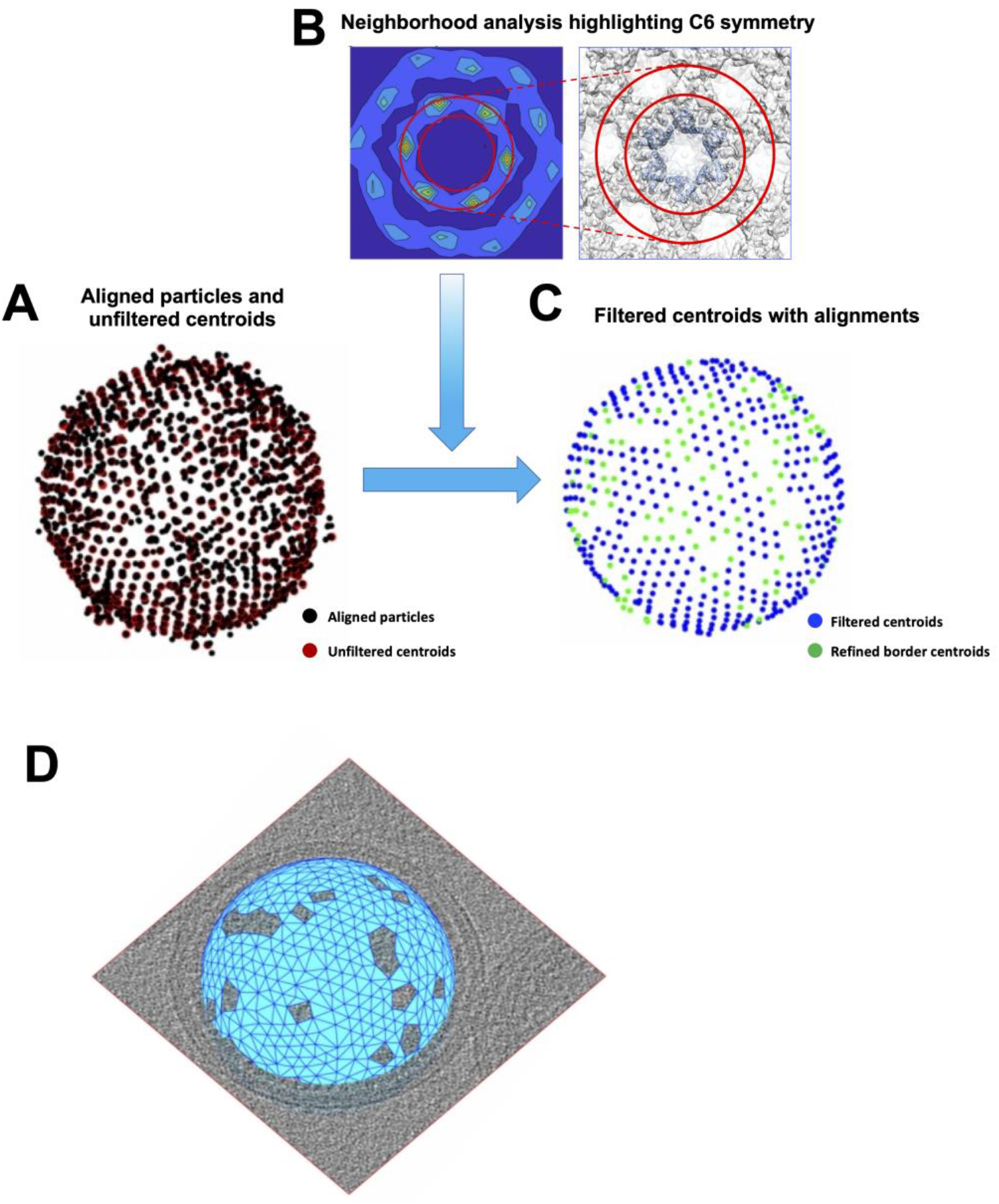
Scheme of the estimation procedure for the Gag layer coverage. (A) Estimated Gag hexamer positions based on the DBSCAN algorithm. A fine initial subtomogram sampling on the expected location of the Gag layer was used in an alignment project with the density map of a single Gag hexamer as a reference, leading to the aligned positions marked as black points. Using the DBSCAN algorithm to identify clusters of putative positions, the initial, unfiltered estimation of Gag hexamer centroids, marked as red points in the panel, are obtained. (B) The neighborhood analysis of unfiltered centroids is a region used for the neighborhood analysis of the initial estimations. The inherent C6 symmetry of the found neighboring Gag hexamer centroids is highlighted. The inner and outer red rings indicate distances of 6.6 and 7.9 nm radii respectively, corresponding to the apparent range of distances between two adjacent Gag hexamer centers. This emergence of the C6 symmetry of the Gag layer is used to filter the initial estimation of the Gag hexamer center positions. (C) Illustration of the Gag hexamer positions determined by the neighborhood analysis algorithm. As a result of this filtering, a subset of the unfiltered centroids was obtained, depicted as blue points, with a local lattice structure around them. This subset was increased by recovering the centroids in positions that follow the C6 symmetry with the filtered centroids. This allowed for refinement of the area on the borders of holes in the Gag lattice. (D) The final depiction of the Gag lattice layer on a HIV-2 immature particle. The blue mesh surface was used to calculate the percentage of the surface area that is covered by Gag hexamers.

**Figure S6.**
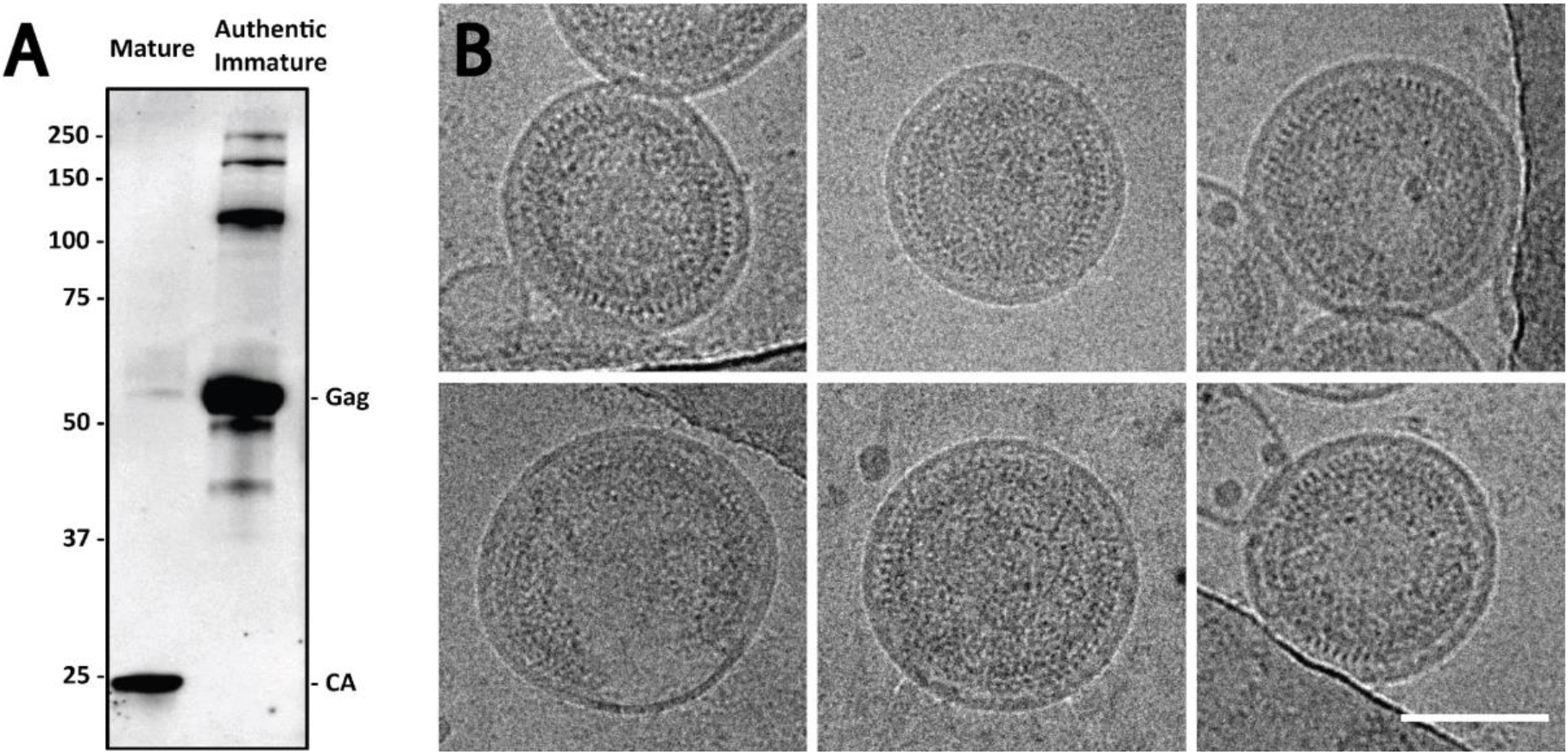
Immunoblot and cryo-EM analysis of authentic immature HIV-2 particles. (A) Immunoblot analysis of mature (WT) and authentic immature (protease catalytic domain mutant) HIV-2 particles using an anti-HIV-2 CA primary antibody which recognizes an epitope within the Gag CA region. (B) Cryo-EM images of authentic immature HIV-2 particles. The scale bar represents 100 nm.

**Figure S7.**
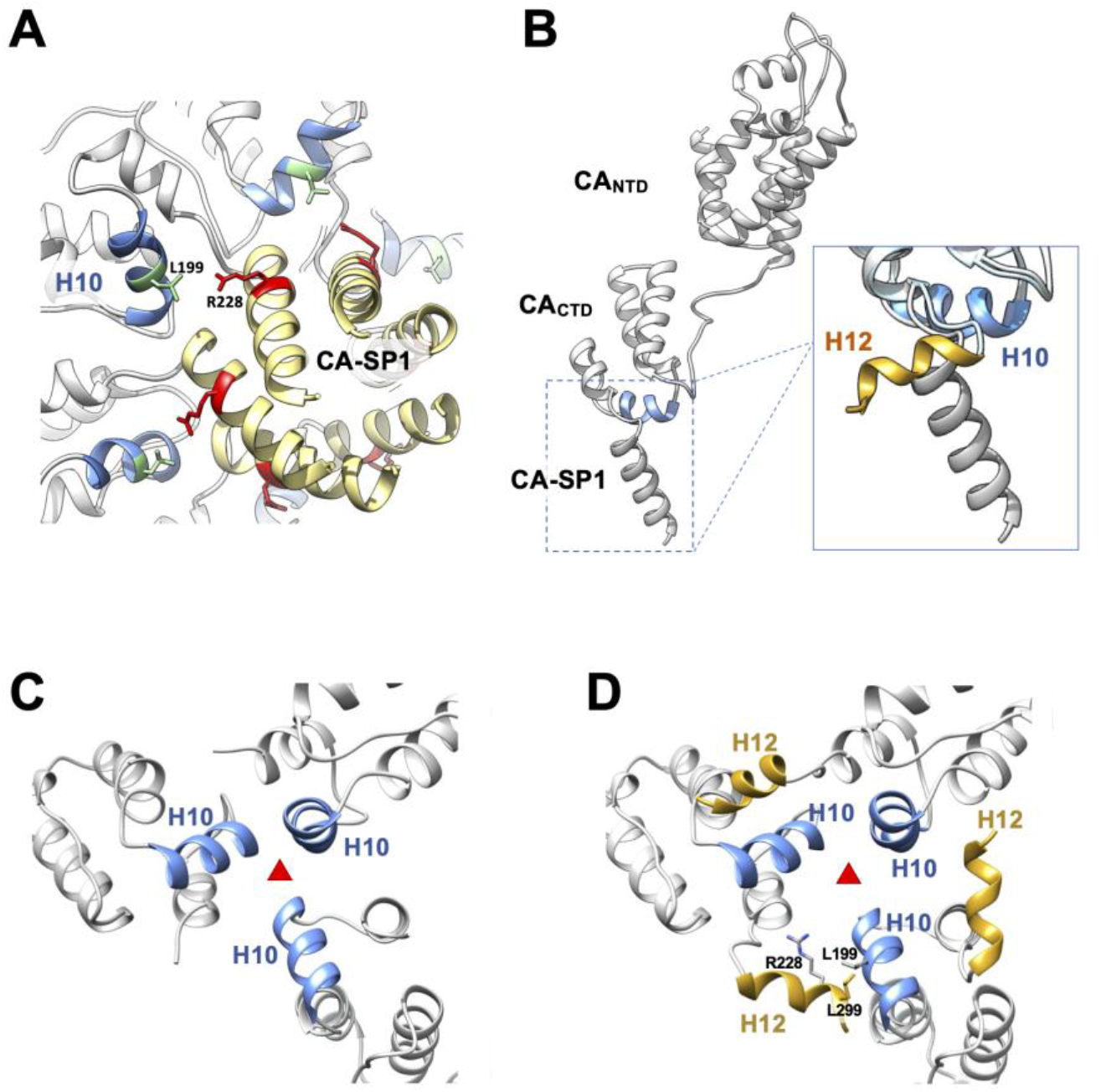
Modeling implications of HIV-2 CA H12 in the immature and mature lattice configuration. (A) Modeled HIV-2 CA and SP1 in the immature Gag lattice, showing CA-SP1 6HB (tan) and H10 (light blue). The residues L199 are colored in light green. The residues of R228 are colored red. (B) Superposition of HIV-2 CA_CTD_ crystal structure on to the HIV-2 immature CA and SP1 model (grey). H10 is colored in light blue, and H12 is colored in gold. Structural comparison shows that H12 of the crystal structure adopts a different conformation of the CA-SP1 helix of the immature CA model. (C) Three-fold interface of mature HIV-1 CA lattice (PDB ID: 6SKN) [71] with H10 indicated in light blue. (D) Three copies of HIV-2 CA_CTD_ fit to HIV-1 CA_CTD_ positions, revealing putative H10-12 interfaces. The red triangle represents the three-fold symmetry axis.

**Figure S8.**
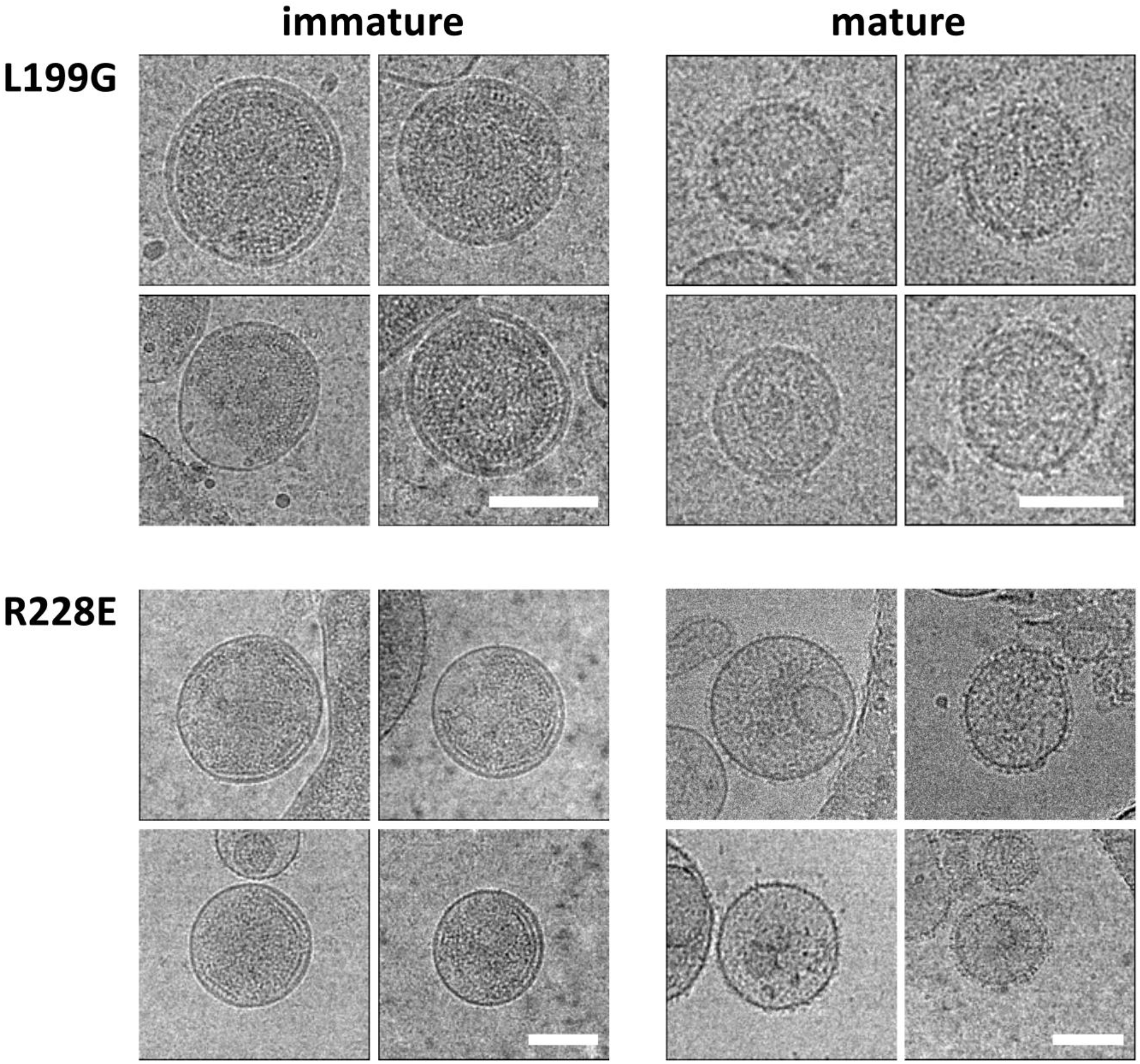
Morphology of HIV-2 Gag L199G and R228E immature and mature particles. Cryo-EM images of the HIV-2 Gag L199G and R228E mutants reveal immature particles with ordered Gag assemblies (left) and the defective CA mature particles with the same mutations in Gag (right). The scale bars represent 100 nm.

**Figure S9.**
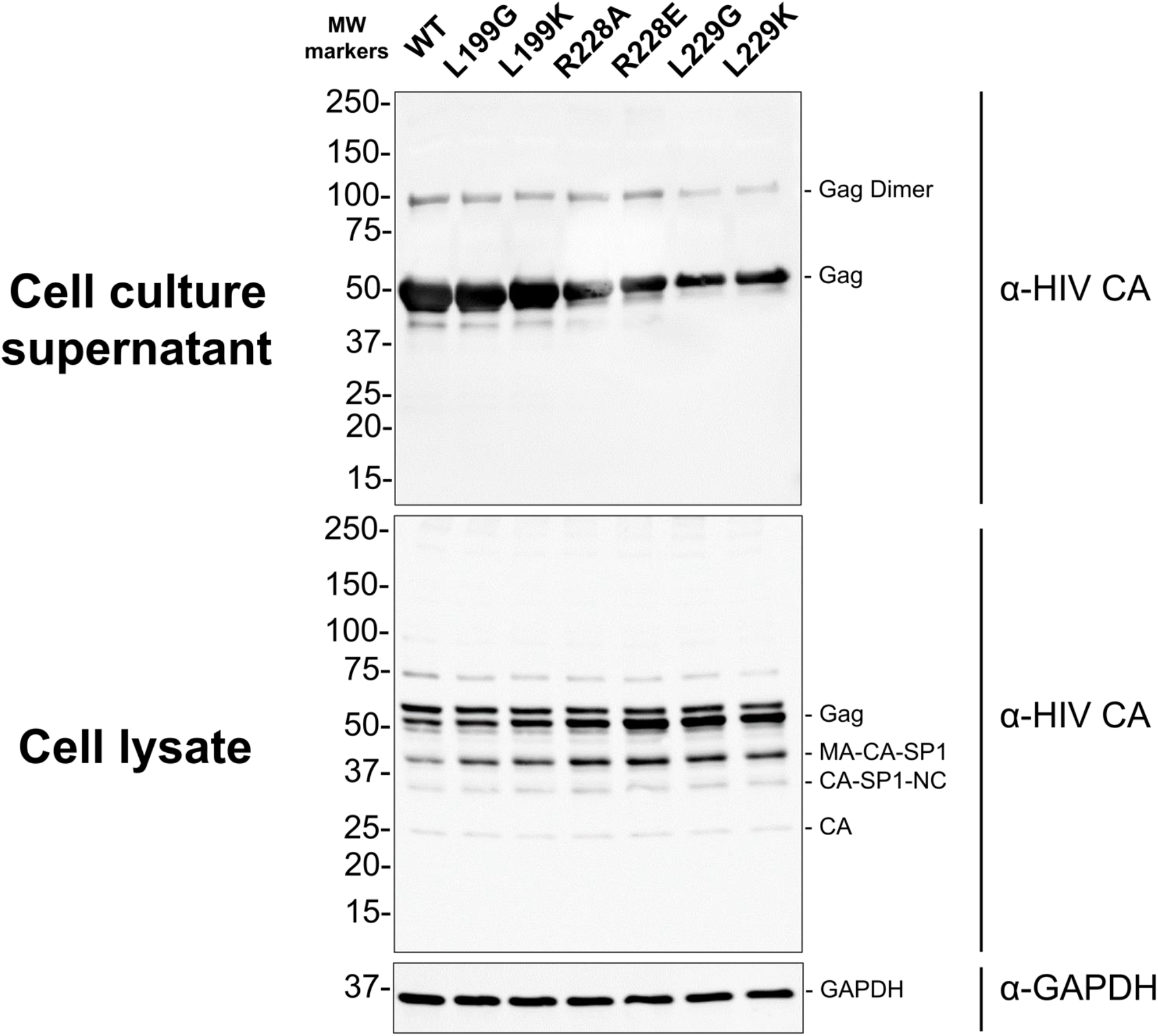
Immunoblot analysis of immature particle production for HIV-2 CA mutants. Immature particle production for HIV-2 CA mutants were analyzed by using an antibody directed against the CA domain of Gag. The Gag proteins from cell culture supernatants were detected with a 1:1500 dilution of anti-HIV-1 p24 antibody in 5% milk TBST and cell lysates were detected with 1:2000 dilution of anti-HIV-1 p24 antibody in 5% milk TBST. Glyceraldehyde 3-phosphate dehydrogenase (GAPDH) in cell lysate was detected by using a 1:10,000 anti-GAPDH antibody in 5% milk TBST. Membranes were washed before incubation with 1:5000 IRDye® 800CW goat anti-mouse IgG secondary antibody and 1:5000 IRDye® 680RD goat anti-rabbit IgG secondary antibody. Immunoblots were imaged by using a ChemiDoc Touch system and analyzed with ImageJ. Gag proteins and Gag cleavage products (i.e., Gag dimer, Gag, MA-CA-SP1, CA-SP1-NC, and CA) and GAPDH (to ensure equal loading of cell lysates) are identified. The sizes of the molecular weight (MW) markers are indicated along the left side of the immunoblot.

**Figure S10.**
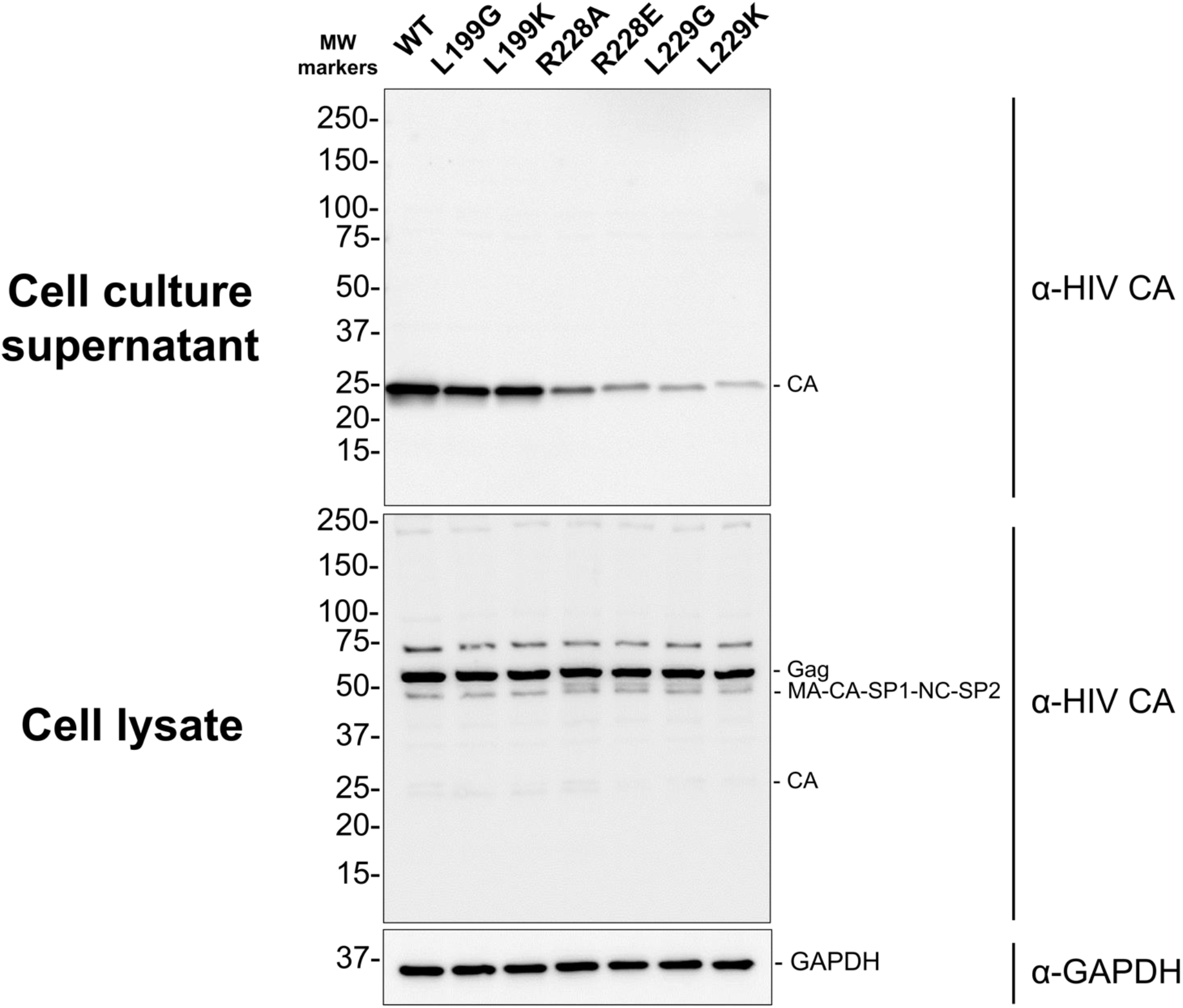
Immunoblot analysis of mature particle production for HIV-2 CA mutants. Mature particle production for HIV-2 CA mutants were analyzed by using an antibody directed against the CA domain. The CA proteins from cell culture supernatants were detected with a 1:1500 dilution of anti-HIV-1 p24 antibody in 5% milk TBST (**Figure 5B**) and cell lysates were detected with 1:2000 dilution of anti-HIV-1 p24 antibody in 5% milk TBST. Glyceraldehyde 3-phosphate dehydrogenase (GAPDH) in cell lysate was detected by using a 1:10,000 anti-GAPDH antibody in 5% milk TBST. Membranes were washed before incubation with a 1:5000 IRDye® 800CW goat anti-mouse IgG secondary antibody and 1:5000 IRDye® 680RD goat anti-rabbit IgG secondary antibody. Immunoblots were imaged by using a ChemiDoc Touch system and analyzed with ImageJ. Gag proteins and Gag cleavage products (i.e., Gag-Pol, Gag, MA-CA-SP1-NC-SP2, and CA) and GAPDH (to ensure equal loading of cell lysates) are identified. The sizes of the molecular weight (MW) markers are indicated along the left side of the immunoblot.

**Figure S11.**
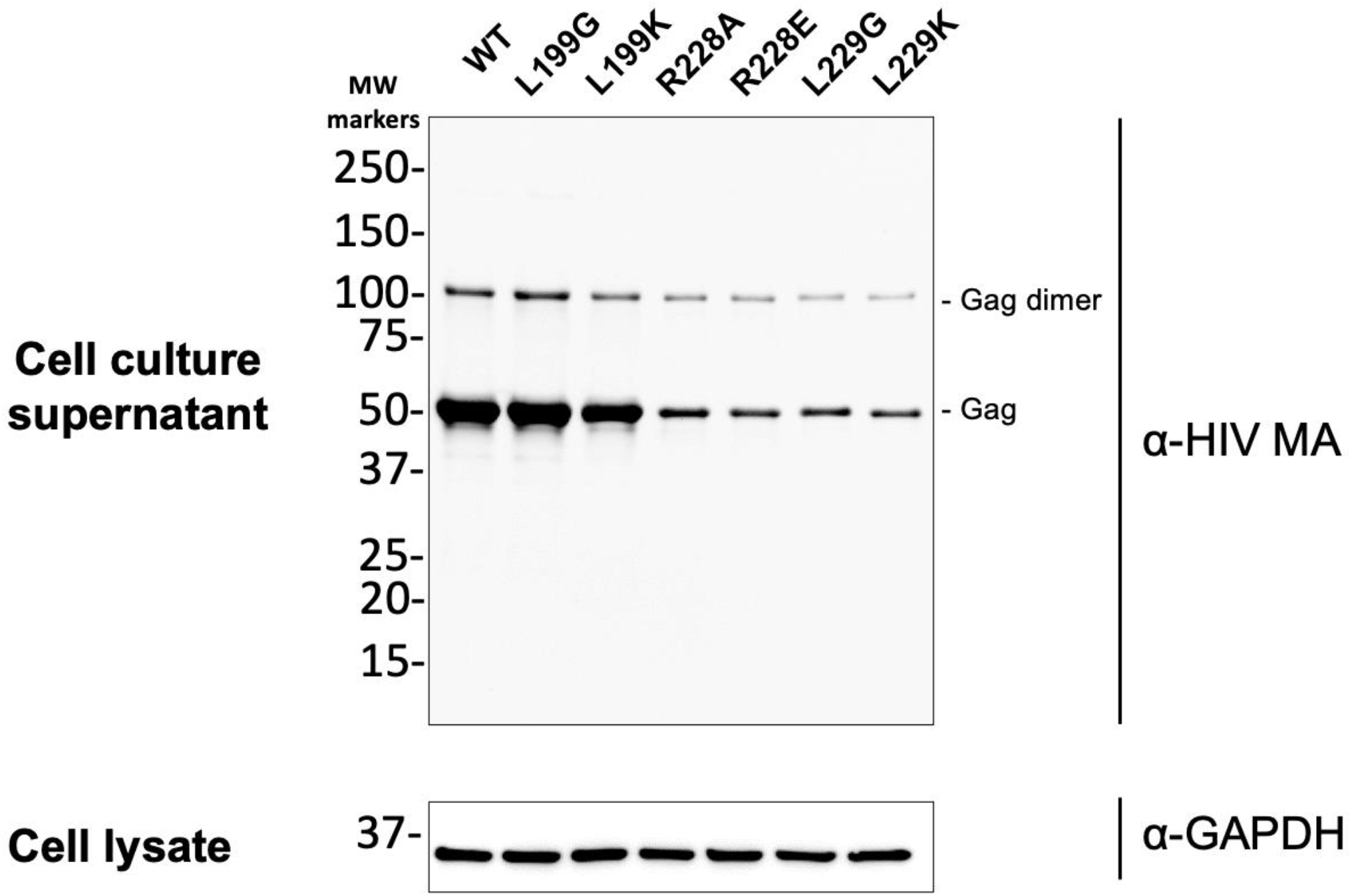
Confirmation of immature particle production for HIV-2 CA mutants by using anti-MA antibody. Immature particle production for HIV-2 CA mutants were analyzed by using an antibody directed against the MA domain of Gag (rather than an anti-CA antibody). Gag protein from culture supernatants were detected with a 1:1000 dilution of anti-HIV-1 p17 antibody in 5% milk TBST. Membranes were washed before incubation with a 1:6000 IRDye® 800CW goat anti-mouse IgG secondary antibody. Immunoblots were imaged by using a ChemiDoc Touch system and analyzed with ImageJ. Shown is a representative immunoblot from cell culture supernatant and the GAPDH from cell lysates to confirm particle production from comparable numbers of cells.

